# Hidden genomic features of an invasive malaria vector, *Anopheles stephensi*

**DOI:** 10.1101/2020.05.24.113019

**Authors:** Mahul Chakraborty, Arunachalam Ramaiah, Adriana Adolfi, Paige Halas, Bhagyashree Kaduskar, Luna Thanh Ngo, Suvratha Jayaprasad, Kiran Paul, Saurabh Whadgar, Subhashini Srinivasan, Suresh Subramani, Ethan Bier, Anthony A. James, J.J. Emerson

**Author notes:** These authors contributed equally to this work (Mahul Chakraborty, Arunachalam Ramaiah). Correspondence to: J.J. Emerson.

## Abstract

**Background:** The mosquito *Anopheles stephensi* is a vector of urban malaria in Asia that recently invaded Africa. Studying the genetic basis of vectorial capacity and engineering genetic interventions are both impeded by limitations of a vector’s genome assembly. The existing assemblies of *An. stephensi* are draft-quality and contain thousands of sequence gaps, potentially missing genetic elements important for its biology and evolution.

**Results:** To access previously intractable genomic regions, we generated a reference-grade genome assembly and full transcript annotations that achieve a new standard for reference genomes of disease vectors. Here, we report novel species-specific transposable element families and insertions in functional genetic elements, demonstrating the widespread role of TEs in genome evolution and phenotypic variation. We discovered 29 previously hidden members of insecticide resistance genes, uncovering new candidate genetic elements for the widespread insecticide resistance observed in *An. stephensi*. We identified 2.4 Mb of the Y-chromosome and seven new male-linked gene candidates, representing the most extensive coverage of the Y-chromosome in any mosquito. By tracking full length mRNA for >15 days following blood feeding, we discover distinct roles of previously uncharacterized genes in blood metabolism and female reproduction. The Y-linked heterochromatin landscape reveals extensive accumulation of long-terminal repeat retrotransposons throughout the evolution and degeneration of this chromosome. Finally, we identify a novel Y-linked putative transcription factor that is expressed constitutively through male development and adulthood, suggesting an important role throughout male development.

**Conclusion:** Collectively, these results and resources underscore the significance of previously hidden genomic elements in the biology of malaria mosquitoes and will accelerate development of genetic control strategies of malaria transmission.

## Background

Mosquitoes transmit the largest number of arthropod vector-borne diseases (i.e. Malaria, Dengue, Zika, Yellow fever and Chikungunya) in humans and animals globally [1, 2]. The complex disease human Malaria is caused by *Plasmodium* parasites, which are transmitted by *Anopheles* mosquitoes [3]. Nearly 20 years ago, sequencing of the *An. gambiae* genome catalyzed rapid growth in genetics and genomics research in this important vector of sub-Saharan Africa [4–6]. However, lack of comparable genomic resources in other malaria vectors have impeded progress in understanding and control of the spread of this deadly disease in other continents [7, 8].

*Anopheles stephensi* is the primary vector of urban malaria in the Indian subcontinent and the Middle East and an emerging malaria vector in Africa [9, 10]. The species is so invasive that without immediate control, it is predicted to become a major urban malaria vector in Africa, putting 126 million urban Africans at risk [11]. Genetic strategies (e.g. Clustered Regularly Interspaced Short Palindromic Repeats (CRISPR) gene drive) that suppress or modify vector populations are powerful means to curb malaria transmission [12, 13]. Success of these strategies depends on the availability of accurate and complete genomic target sequences and variants segregating within them [12, 14]. However, functionally-important genetic elements and variants within them often consist of repetitive sequences that are either mis-assembled or completely missed in draft-quality genome assemblies [15, 16]. Despite being a pioneering model for transgenics and CRISPR gene drive in malaria vectors [13, 17], the community studying *An. stephensi* still relies on draft genome assemblies that do not achieve the completeness and contiguity of reference-grade genomes [7, 18]. This limitation obscures genes and repetitive genetic elements that are potentially relevant for understanding parasite transmission or for managing vector populations [19]. We therefore generated a high-quality reference genome for a laboratory strain UCISS2018 (see Methods) of this mosquito sampled from the Indian subcontinent (fig. S1) using deep coverage long reads plus Hi-C scaffolding and then annotated it by full length, mRNA sequencing (Iso-Seq). These resources facilitate characterization of regions of the genome less accessible to previous efforts, including gene families associated with insecticide resistance, targets for gene-drive interventions, and recalcitrant regions of the genome rich in repeats, including the Y chromosome.

## Results

### A reference-grade assembly of *An. stephensi*

*Anopheles stephensi* has three major gene-rich chromosomes (X, 2, 3) and a gene poor, heterochromatic Y chromosome (Fig. 1A) [18]. The size of the published most contiguous draft assembly of *An. stephensi* genome was 221 Mb and had 23,371 scaffolds with N50 of 1.59 Mb, meaning that half of the assembly is found in scaffolds <1.59 Mb (table S1) [18]. The longest scaffold of this assembly was 5.9 Mb and 11.8 Mb of gaps in the assembly were filled with Ns. In the new reference assembly, the major chromosomes are represented by just three sequences (scaffold N50 = 88.7 Mb; contig N50 = 38 Mb; N50 = 50% of the genome is contained within sequence of this length or longer), making this assembly comparable to the *Drosophila melanogaster* reference assembly, widely considered a gold standard for metazoan genome assembly [20] (Fig. 1B, Table 1, figs. S4-5, table S1, supplementary text). The new reference assembly has 89% (205/235 Mb) of the estimated *An. stephensi* physical haploid genome assigned to chromosomes, which parallels the assembly completeness of *An. gambiae* where 88% (230/265Mb of physical genome size) of the assembled genome is placed into chromosomes (see Methods) (Fig. 1).

**Fig. 1.**
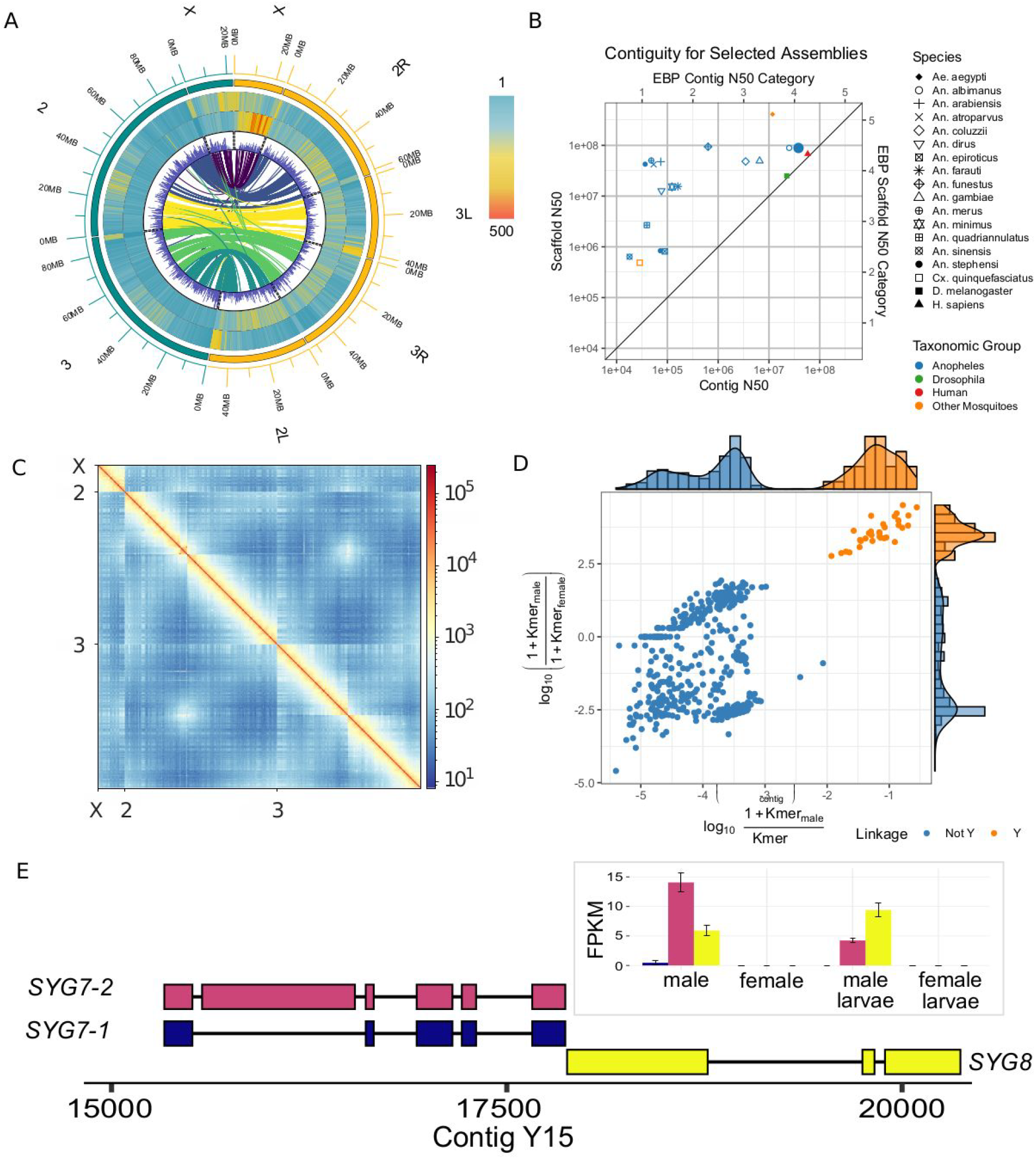
*Anopheles stephensi* genome assembly. (**A**) Distribution of repeats, gene content, and synteny between *An. stephensi* (left, green) and *An. gambiae* (right, yellow) genomes. Each successive track from outside to inside represents TE density, satellite density (fig. S2, fig. S3), and gene density across the chromosome arms in 500 kb windows. The innermost track describes the syntenic relationship between *An. stephensi* and *An. gambiae* chromosome arms. (**B**) Contiguities of published genome assemblies of *Anopheles* malaria vectors, *Culex*, *Aedes aegypti*, human (GRCh38.p13) and the model organism *D. melanogaster*. Among the mosquito vectors, *An. stephensi* assembly reported in the current study is the only genome that matches the Earth BioGenome (EBP) standard of the human and the Drosophila genomes [25]. (**C**) Hi-C contact map of the *An. stephensi* scaffolds. Density of Hi-C contacts are highest at the diagonals, suggesting consistency between assembly and the Hi-C map. (**D**) Identification of putative Y contigs using the density of male-specific k-mers on the x-axis and the ratio of male and female k-mers on the y-axis. (**E**) Transcripts of *SYG7* and *SYG8*, two new Y-linked genes as revealed by the uniquely mapping Iso-seq reads. SYG7 has two isoforms. Inset: transcript abundance of *SYG7* and *SYG8* in male and female adults and larvae. As shown here, neither gene is expressed in females.

**Table 1.** Summary of genome assembly and annotation statistics.

| Genomic features | Value |
| --- | --- |
| Total length (bp) | 250,632,892 |
| Contig number | 566 |
| Contig N50 (bp) | 38,117,870 |
| Scaffold Number | 560<sup>^</sup> |
| Scaffold N50 (bp) | 88,747,609<sup>*</sup> |
| L50 | 2 |
| GC content (%) | 44.91% |
| Maximum scaffold length | 93,706,523 |
| Minimum scaffold length | 1727 |
| N's per 100 Kb | 3.60 |
| Alternative haplotypes, bp (# scaffolds) | 15,743,318 (169) |
| Unclassified contigs, bp (# scaffolds) | 20,118,109 (355) |
| Putative Y-chromosome, bp (# scaffolds) | 2,431,719 (33) |
| 3 major chromosomes X, 2 & 3, bp (# scaffolds) | 205,167,748 (3) |
| Predicted Genes | 14,966 |
| Predicted Transcripts | 16,559 |
| 5' UTR | 9,791 |
| 3' UTR | 9,290 |
| tRNAs | 503 |

<sup>^</sup>Except three major chromosomes, we kept others as contigs; <sup>\*</sup>Scaffold N50 is the length of chr3.

The new *An. stephensi* assembly recovers 99.2% of 3,285 complete single copy Diptera orthologs (i.e. Benchmarking Universal Single Copy Orthologs or BUSCOs). The reference assembly of the *D. melanogaster* genome captures 99.1% of BUSCOs, indicating that their completeness is comparable (Table 1, fig. S5, Supplementary text). The *An. stephensi* assembly not only achieves dramatic improvements over the existing draft assembly (1,044-fold and 56-fold increase in contig N50 and scaffold N50, respectively and a 97% reduction in assembly gaps), but it is also more contiguous and complete than the latest version of published assemblies of *Anopheles* species (single copy BUSCOs 93.7-98.9%), including extensively studied African malaria vector *An. gambiae* (AgamP4) genome (BUSCO 97.4%, contig N50 = 85 kb, scaffold N50 = 49 Mb), arguably the best characterized genome among malaria vectors (Table 1, table S1, Fig. 1A and B, fig. S3, fig. S5) [7, 19, 21–24]. Furthermore, the concordance of short reads mapped to the assembly suggests that assembly errors are rare (short read consensus quality value or QV = 49.2, or ~1 discrepancy per 83 kb), which is further supported by uniformly mapping long reads and a high resolution Hi-C contact map (Fig. 1C, fig. S1, Table 1). Among the mosquito vectors, the *An. stephensi* assembly reported in here is the genome that most closely matches the standards achieved by the human and the *Drosophila* genomes (Fig. 1B) and is the only *Anopheles* genome to meet the standards of Earth BioGenome Project (EBP) standard [25] for both contiguity and accuracy.

Further, assessment of annotation showed that 92% of genes annotated with <0.5 annotation edit distance (AED), which surpasses the recommended score (>90%) for gold standard annotations (Campbell et al. 2014). The Pfam content score (65.4%) also above the recommended range 55-65%, indicating that the *An. stephensi* proteome is well annotated. Comparison of gene model annotations of our assembly with the draft assembly showed that 24.4% of 14,966 genes were unique in our assembly. An additional 1,429 genes in our assembly were split over >1 contigs in the older most contiguous *An. stephensi* assembly [18]. Collectively, such evidence suggests that our assembly recovered previously unassembled functional genome. We also assembled 33 putative Y contigs totaling 2.4 Mb, representing the most extensive Y chromosome sequence yet recovered in any *Anopheles* species [26] (Fig. 1D to F) [7, 18, 26]. Finally, to assist disease interventions using endosymbionts [27], we assembled *de novo* the first complete genome of the facultative endosymbiont *Serratia marcescens* from *Anopheles* using sequences identified in the *An. stephensi* long read data (fig. S4).

### Transposable elements

As naturally-occurring driving genetic elements, transposable elements (TEs) are invaluable for synthetic drives [28, 29] and transgenic tools [17, 30, 31]. Although TEs comprise 11% (22.5 Mb) of the scaffolded *An. stephensi* genome, the most contiguous published draft assembly [18] contains 32% or 7.2 Mb less TE sequences, the majority of which (61% or 4.4 Mb) are composed of previously unknown LTR retrotransposons (Fig. 2A). The proportion of absent TE sequences in the draft assembly resembles the share of TE sequences missed by short reads based TE detection [15]. Most full length LTR and non-LTR retrotransposons we identified were either absent or fragmented in the existing draft genome assembly (Fig. 2A, fig. S2C, table S2). Some of these TEs are likely strain-specific and are absent from the strain sequenced previously [18]. The newly identified TEs include species-specific and evolutionarily recent retrotransposons, which highlight the dynamic landscape of new TEs in this species and provide a resource for modeling the spread of synthetic drive elements (Fig. 2B). The *An. stephensi* genome possesses fewer TEs and satellites than *An. gambiae*, partly accounting for the difference in their genome size and composition of the pericentric regions (Fig. 1A, fig. S2 and fig. S3). The difference in repeat density between the two species is particularly prominent on the X chromosome, which also carries disproportionately higher abundance of TEs and satellites among the three major chromosomes (p < 2.2e-16, proportion test for equal TE content) (Fig. 1A, fig. S2). Although most (98% of 24.8 Mb) TEs are located within introns and intergenic sequences [32], we observed 1,368 TE sequences in transcripts of 8,381 Iso-Seq supported genes, 68% (939/1,368) of which were not found in the earlier assembly [18] (table S3).

**Fig. 2.**
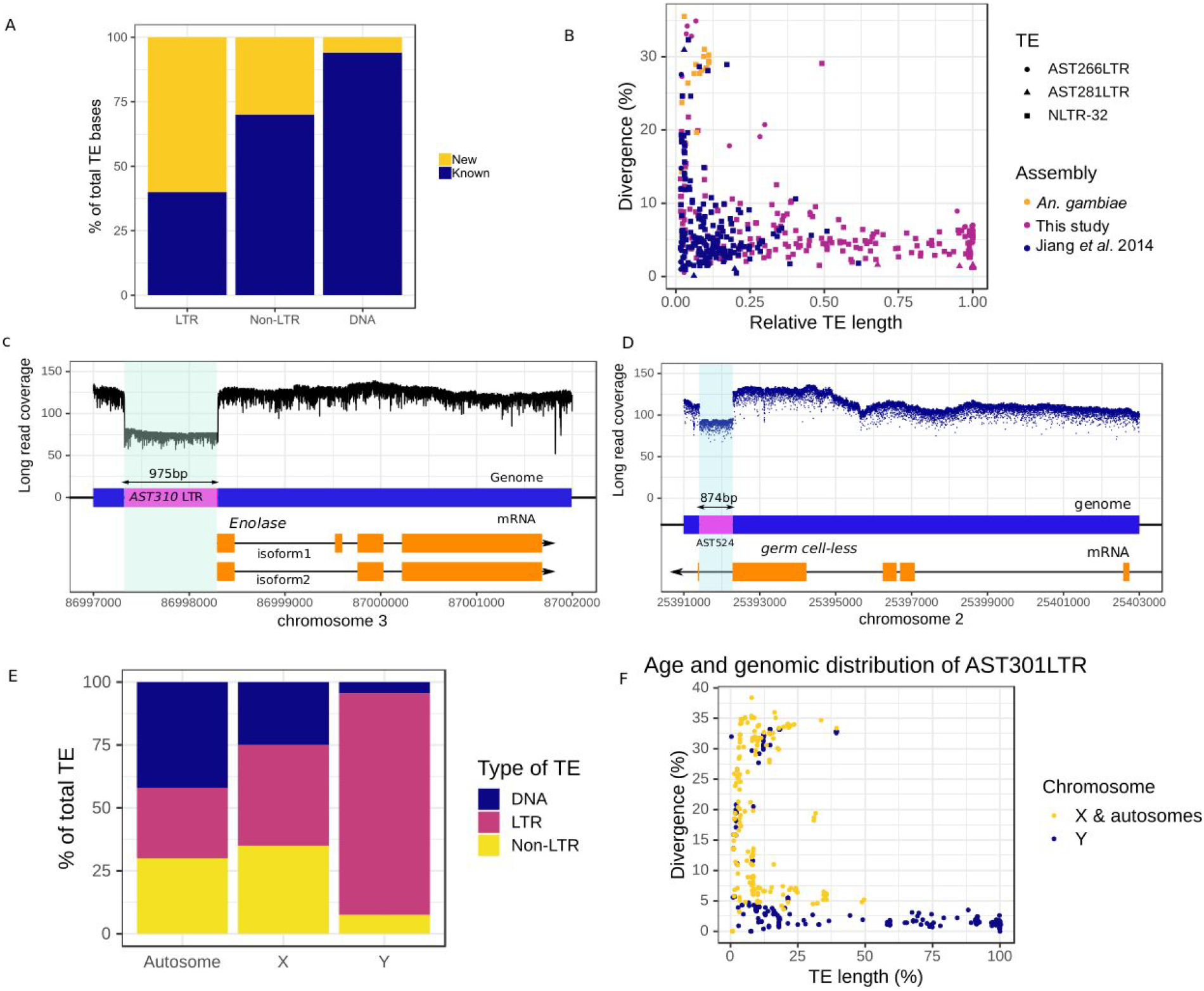
TEs and their role in genetic variation in*An. stephensi.* (**A**) Proportion of TEs (counted in bp) that were uncovered by the new reference assembly of *An. stephensi*. Many LTR and non-LTR TEs are identified for the first time. (**B**) Similar to many other LTR and non-LTR retrotranspososons, LTR elements AST266LTR and AST281LTR and non-LTR element NLTR-32 do not have any closely-related counterpart in *An. gambiae*. As shown here, only small parts of these TE sequences were known. (**C**) Insertion of a polymorphic LTR fragment immediately upstream of the highly-conserved gene *Enolase*.The insertion creates a gap between the promoters and the transcription start site in half of the alleles and may disrupt transcription of the gene. (**D**) A polymorphic DNA element AST524 located in the 3’ UTR of *gcl* creates a null *gcl* allele. (**E**) Comparison of TE compositions in the autosomal, X and Y chromosomal sequences. Not only are most of Y sequences repeats (fig. S6), but the majority of Y TEs are LTR elements. (**F**) LTR retrotransposon AST301 is present in intact copies only, in Y contigs. Its counterparts in the autosomes and X chromosome are fragmented and more diverged than the AST301 sequences found on the Y sequences.

Due to the low, but measurable, level of residual heterozygosity in the sequenced, strain (fig. S1; see Methods), we discovered several segregating TEs (table S4, fig. S6), some of which likely have functional consequences. For example, a 975 bp LTR fragment inserted immediately at the 5’ end of the *Enolase* gene (*Eno*) may perturb its transcription (Fig. 2C). In *D. melanogaster*, null mutants of *Eno* show severe fitness and phenotypic defects that range from flightlessness to lethality [33], whereas reduction in *Eno* expression protects *Drosophila* from cadmium and lead toxicity [34]. Because *Eno* is highly conserved between *An. stephensi* and *D. melanogaster* (88% of 441 amino acids are identical between the two), we anticipate that this structural variant (SV) allele of *Eno* might be deleterious, although it also could confer some degree of resistance to heavy metal toxins. Another TE, a 874 bp DNA element, is inserted into the 3’ UTR of the gene, *germ cell-less* (*gcl*), the *Drosophila* ortholog of which determines germ cell development [35] (Fig. 2D). All full-length Iso-Seq reads from this gene are from the non-insertion allele, suggesting that the insertion allele is a *gcl* null.

### Structural features and newly discovered genes of the Y chromosome

Targeting Y-linked sequences can be the basis for suppressing vector populations [36]. We identified 33 putative Y contigs we identified using a k-mer based approach (Fig. 1D). We experimentally validated three unique sequences spread across three contigs using PCR, which confirmed the male specificity of the predicted Y sequences (see Methods). In contrast to the autosomes and the X chromosome, 72% of the Y sequences (1.7 Mb) comprise LTR elements (Fig. 2E, fig. S6). While most full length LTR elements in the Y chromosome also are present in the other chromosomes, a 3.7 kb retrotransposon, AST301, is represented by 46 highly-similar (<2.5% divergent) full-length copies only in the Y sequences. Its matching sequences elsewhere in the genome are small and evolutionary distant, consistent with AST301 being primarily active in the Y chromosome (Fig. 2F). The proliferation of AST301 may be a consequence of the Y chromosome’s lack of recombination, which can lead to irreversible acquisition of deleterious mutations in a process called Muller’s ratchet [37].

Despite the high repeat content, we uncovered seven Y-linked genes that are supported by multiple uniquely mapping Iso-Seq reads (table S5, fig. S7). We also recovered the three previously-identified Y-linked genes and filled sequence gaps in *Syg1* [38]. Two of the newly-discovered Y-linked genes (*Syg7* and *Syg8*) sit in a cluster of three overlapping Y-linked genes, all of which show strong expression in male larvae and adults but no expression in larval or adult females (Fig. 1, E and F). Y-linkage of these genes are also confirmed by PCR (see Methods). Both genes show low or absent expression in the early (0-2 hours) embryos but are expressed in the later stages (>4 hours) (fig. S7). Translation of open reading frames from *Syg7* transcripts shows the presence of a myb/SANT-like domain in Adf-1 (MADF) domain in the encoded protein (fig. S7).

### Transcriptional response to blood feeding

An alternative to suppression schemes aimed at reducing mosquito numbers is modification of mosquito populations to prevent them from serving as parasite vectors. Promoters induced in females by blood feeding can be repurposed to express effector molecules that impede malaria parasite transmission [39, 40]. Following a blood meal, hundreds of genes are induced, many of which stay upregulated for days after the blood meal (Fig. 3A, table S6). When ranked by expression-fold changes post blood meal (PBM), the top 1% of genes with most strongly affected expression (representing a >64-fold change) include 593 genes enriched for involvement in DNA replication, cell division, amino acid metabolism, and signalling within and between cells (Fig. 3B). Comparison of protein sequences of the upregulated genes and their orthologs in *An. gambiae* suggest that most genes are relatively conserved (70% share >80% identity) (Fig. 3C). This suggests that the genes likely retained similar functions in the two species. However, sequences of 19.3% (115/593) of these genes are either fragmented or unannotated in the draft *An. stephensi* assembly [18] (table S7). For example, the eye pigmentation gene *white*, a key genetic marker in a wide range of insects and showing upregulation following a blood meal, was fragmented in the draft assembly due to the presence of an 853 bp intronic non-LTR retrotransposon (Fig. 3D). Similarly, despite the active roles of blood-inducible and constitutively-expressed protease genes encoding trypsins and chymotrypsins in digestion and metabolism of blood [41, 42], not all of their complete sequences in *An. stephensi* were known before this study (table S7).

**Fig. 3.**
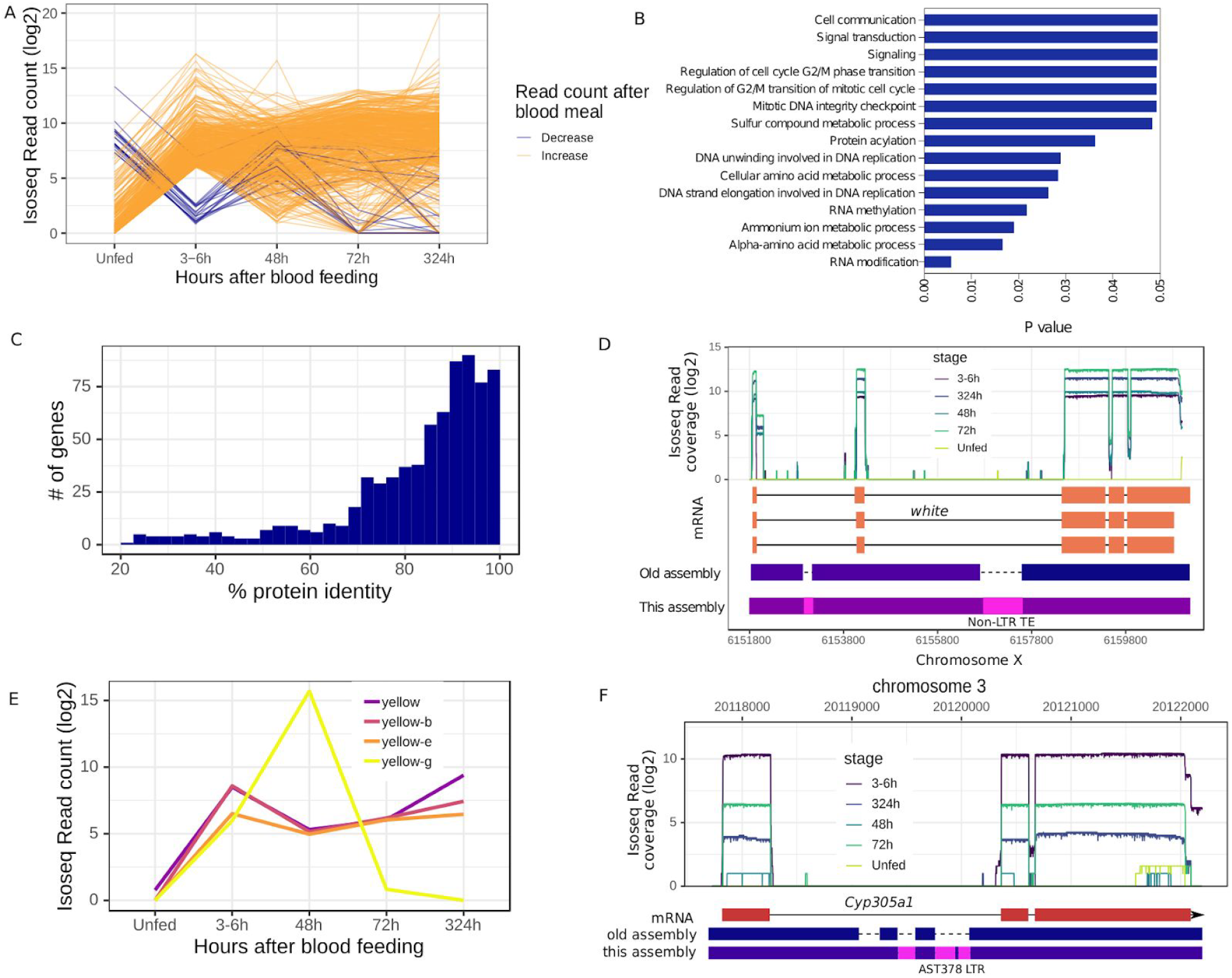
Gene expression changes in adult female mosquitoes after a blood meal. (**A**) Transcript abundance of genes that are in the top 1% (>~64 fold) of the PBM transcript abundance changes. As evident here, more genes show upregulation than downregulation, although expression changes of some genes may not be due to the blood meal. (**B**) GO gene enrichment analysis of the genes from panel A. Consistent with the role of the blood meal in mosquito biology, the genes involved in cell division, DNA replication, amino acid metabolism, and cell signalling are enriched among the differentially-expressed genes. (**C**) Protein sequence identity between the *An. stephensi* genes showing PBM upregulation and their *An. gambiae* orthologs. (**D**) Despite being a common genetic marker, sequence of the PBM upregulated *white* gene was fragmented in the draft assembly of *An. stephensi*. (**E**) Transcript abundance of four *yellow* genes (*yellow*, *yellow-b*, *yellow-e*, *yellow-g*) before and after blood meal. All genes show a similar transcript profile until 6 hours PBM after which *yellow-g* transcripts become more abundant. (**F**) A *Cyp450* orthologous to *D. melanogaster Cyp305a1* shows PBM upregulation and harbors intronic TEs are absent in the Jiang et al. (2014) assembly.

A Yellow protein gene (*yellow-g*) that was shown previously to be essential for female reproduction in *An. gambiae* and was used as a target in a CRISPR gene drive [43] was found to be upregulated PBM. Yet, neither *yellow-g* nor the three other members of the Yellow protein gene family (*yellow-b*, *yellow-e*, *yellow*) that showed PBM elevated transcript levels were previously annotated (Fig. 3E) [42]. Although transcript levels of all four genes increase in tandem until 6 hours PBM, the *yellow-g* transcript level continues to climb until 48 hours PBM and then reverts to the pre-BM level (Fig. 3E). In contrast, the other three *yellow* genes maintained a similar transcript level even after 13 days (Fig. 3E). The PBM upregulation pattern of the four yellow genes in *An. stephensi* is consistent with -roles in female reproduction– although *yellow-b*, *yellow-e*, and *yellow* are probably required longer than *yellow-g*. Interestingly, a Cytochrome P450 monooxygenase (Cyp450) gene, which bears similarity to *D. melanogaster Cyp305a1*, also was upregulated (Fig. 3F). *Cyp305a1* acts as an epoxidase in the Juvenile hormone biosynthesis pathway and helps maintain intestinal progenitor cells in *D. melanogaster* [44]. PBM upregulation of *Cyp305a1* suggests that it may have a potential role in cellular homeostasis in the midgut after a blood meal [45].

The *cis*-regulatory elements of the blood-meal-inducible genes can be combined with antimicrobial peptide genes to explore new effector molecule candidates to block malaria parasite transmission [39]. The new assembly revealed 361 immune-related genes, 103 of which were either previously unknown or broken in the previous assembly [18]. Among the immune-related genes, 15 genes were upregulated and are among the top 1% PBM upregulated genes (table S6). The immunity genes also included 20 putative antimicrobial peptides (AMP), three of which (GENE_00013347, GENE_00002217, GENE_00011093) were not known before (supplementary text, fig. S8, table S8). GENE_00013347 encodes for *Cecropin-A*, ortholog of which in *An. gambiae* protects against infections to gram negative and gram positive bacteria, fungi, and yeasts [46]. The AMPs and other immunity genes we identified provide a rich arsenal of effector molecule candidates for transgenic intervention of *Plasmodium* transmission in *An. stephensi*.

### Insecticide resistance genes

In Asia and eastern Africa, *An. stephensi* populations show widespread resistance to dieldrin, DDT, malathion and pyrethroids [47–49]. Insecticide resistance in these populations has been attributed to various Cytochrome p450s, Esterases, GABA receptor (*Resistance to dieldrin* or *rdl*), and voltage gated sodium channel (*Knock-down resistance* or *kdr*) [50]. Frequently, amino acid changes in *rdl* and *kdr* and copy number increases of Cyp450s, Esterase (Est) and Glutathione S-transferases (*Gst*) have been associated with the resistance phenotypes [48, 50].

We identified 94 Cyp450, 29 *Gst*, and 16 esterase genes including 2 Acetylcholine esterases (*ace-1* and *ace-2*), providing a comprehensive resource for discovery and delineation of the molecular basis of insecticide resistance (see Methods). Sequences of 22% (31/139) of these genes were either fragmented or missing in the draft assembly (see Methods) (Fig. 4, A and B) [18]. We also recovered the complete *kdr* sequence and discovered 8 transcript isoforms for this gene (fig. S9). A polymorphic TE insertion immediately downstream of *kdr* suggests that TE insertions play an important role in genetic variation in insecticide resistance candidate genes. We also resolved tandem arrays of insecticide resistance genes, as evidenced by a 28 kb region consisting of Cyp450s similar to *D. melanogaster Cyp6a14*, *Cyp6a23*, *Cyp6a8*, *Cyp6a18* and *Musca domestica Cyp6A1* (Fig. 4B). In *D. melanogaster*, *Cyp6a14* is a candidate gene for DDT resistance [51], suggesting a similar function for its *An. stephensi* counterpart. One *Cyp6a14* in the array has a polymorphic 191 bp LTR TE fragment inserted 1 kb to the 5’ end of the transcription start site, implying the presence of more than one SV allele in this complex region (Fig. 4C). We also resolved previously fragmented tandem copies of *Esterase B1* (*Est-B1a* and *Est-B1b*) counterparts, which have been shown previously in *Culex quinquefasciatus* to provide resistance to organophosphates [52] (Fig. 4D). Interestingly, the Cyp450s in the array and *Est-B1b* show opposite sex-biased expression, suggesting that the molecular basis of insecticide resistance may differ between sexes (Fig. 4D).

**Fig. 4.**
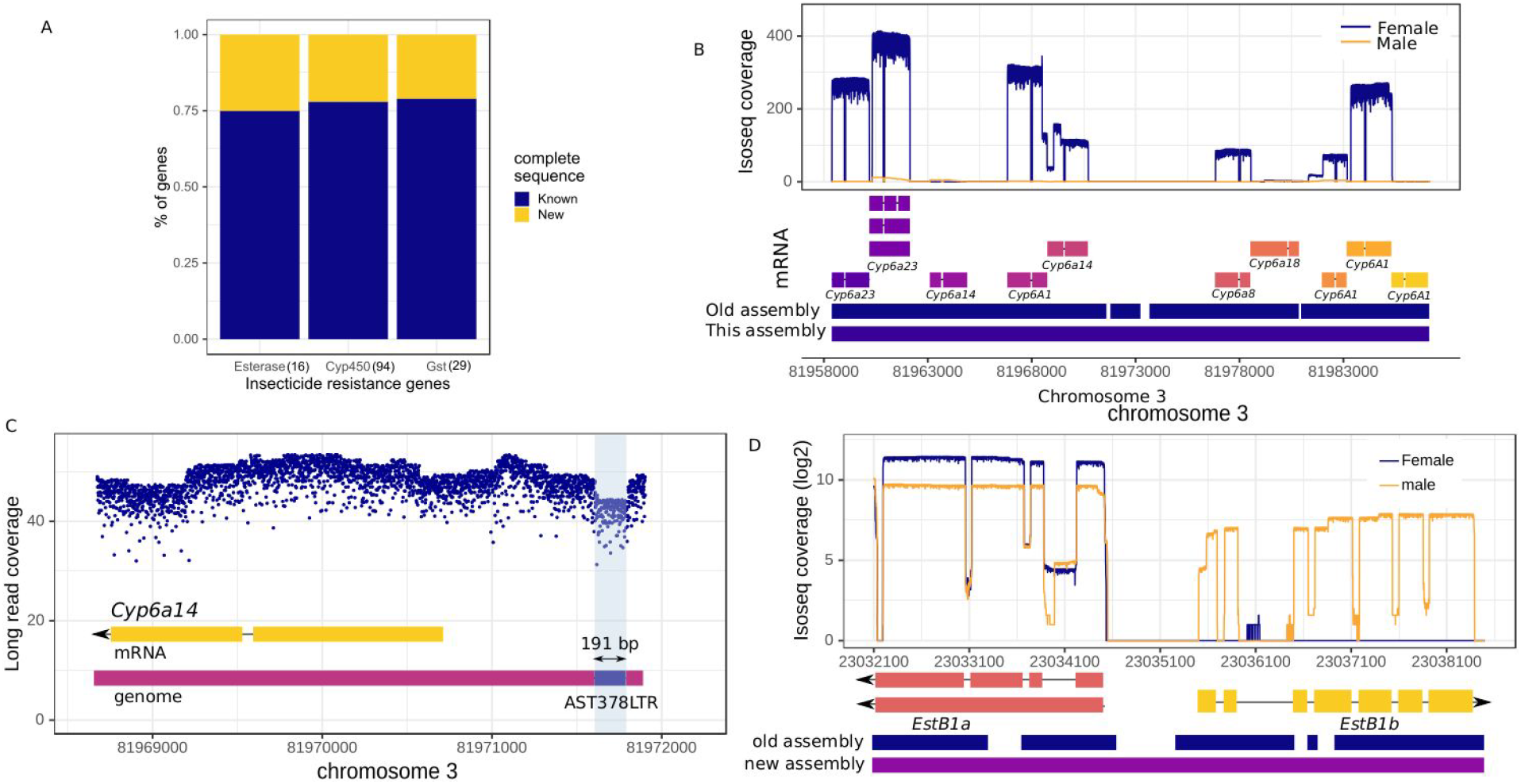
Putative insecticide resistance genes in *An. stephensi*. (**A**) Proportion of various candidate insecticide-resistance genes that are either fragmented or missing repetitive regions in the draft *An. stephensi* assembly. (**B**) An array of tandemly-located *Cyp450* genes that include *Cyp6a14*, a candidate gene for DDT resistance in *D. melanogaster* [51]. In the earlier assembly, this cluster was broken into three sequences, undermining investigation of the functional effects of these genes. Most genes show female-biased expression. (**C**) A polymorphic AST378 LTR fragment insertion is segregating inside the *Cyp450* array shown in B, suggesting presence of more than one SV allele in this genomic region. (**D**) Tandemly located *Esterase B1* genes show different sex-biased expression patterns. *EstB1a* do not show any strong bias towards either sex, whereas *EstB1b* shows male-biased expression. *EstB1* amplification causes organophosphate resistance in *Culex.* The *EstB1a* and *EstB1b* sequences were broken into five pieces in the earlier *An. stephensi* assembly.

## Discussion

Despite being an important malaria vector in Asia and an emerging vector in Africa, the existing genome assemblies of *An. stephensi* remains fragmented and incomplete. We improved the genetic resources for this species by assembling a highly contiguous *de novo* reference genome that recovers sequences relevant to the biology and evolution of the species missing in previous drafts. We found that the genome of *An. stephensi* is actively shaped by species-specific TEs, which likely comprises a major source of genetic variation. Given the generally known strongly deleterious effects of TE insertions [53, 54], the presence of potentially functionally relevant polymorphic TE insertions even in an inbred laboratory strain indicate that individually rare TE mutations could play a major role in the variation in phenotype and fitness of *An. stephensi* natural populations.

Even though the Y chromosome plays an important role in male biology and sexual conflicts, its highly repetitive nature poses steep challenges for assembly and molecular characterization of the Y chromosome in *Anopheles* [7, 8, 18, 26, 38, 55]. Due to the improved Y representation in our assembly, we characterized the repeat and gene content that previously evaded scrutiny. The enrichment and persistence of full length LTR retrotransposons on this chromosome relative to the autosome and the X chromosome indicates an important role of these TEs in degeneration and heterochromatinization of the Y [18]. Moreover, we annotated at least seven previously undiscovered genes on the *An. stephensi* Y chromosome, suggesting that the paucity of Y-linked genes described in the *Anopheles* genus is due in part to technical limitations rather than only the degeneration of the Y. While the biological functions of these genes remain unknown, one (*Syg7*) contains a MADF DNA binding domain found in certain *D. melanogaster* transcription factors [56], suggesting it is a male-specific transcription factor. Thus, the Y sequences we uncovered provide insights into the contribution of Y towards the differences between the two sexes in *An. stephensi*.

In female *Anopheles* mosquitoes, blood meals initiate a cascade of physiological and molecular events involving hormone and signalling pathways, protein digestion and metabolism, gut homeostasis, and egg development. Our results uncovered several genes that show distinct patterns of regulation post blood meal, underscoring their respective roles in these processes. For example, persistence of *yellow-g* transcript for days after a blood meal is consistent with its role in eggshell integrity in both *Drosophila* and *An. gambiae* [57, 58]. On the other hand, Cytochrome P450s like *Cyp305a1* plays a role in hormone signalling and mediates intestinal homeostasis that are necessary for reproduction [59]. Thus, the new assembly combined with the tracking of full length transcripts revealed the components of the complex biological network that are activated by a blood meal in this species. Further experiments using replicated transcript data will provide a comprehensive view of the effect of blood meal on *An. stephensi* biology.

The spread of insecticide resistance in Asian and African *An. stephensi* populations have made identification of insecticide-resistant mutations an urgent priority [47, 50, 60, 61]. We have uncovered TE insertions in candidate insecticide resistance gene *kdr* which is generally investigated only for amino acid variants [62, 63]. TE insertions in the vicinity of a gene can influence its expression due to its epigenetic silencing effect on the expression of the nearby genes [64]. The TE insertion at *kdr* could potentially affect *kdr* expression, contributing to functional variation *at* kdr that would go unnoticed by studies focusing on amino acid variation. Additionally, as we have shown here, candidate genes for insecticide resistance are often present in clusters of tandem copies that are difficult to resolve without a contiguous and error-free assembly. These regions are also segregating for repetitive structural variants (SV), indicating that SVs in repetitive genomic regions could contribute to functional genetic variation implicated in insecticide resistance in *An. stephensi* [65]. Evidently, such assemblies would be key to the detection of causal SV mutations for insecticide-resistance [15, 19].

## Conclusion

CRISPR and gene drive-based strategies promise to transform the management of disease vectors and pest populations [66]. However, safety and effectiveness of these approaches rely on an accurate description of the functional and fitness effects of the genomic sequences and their variants [12, 67]. Draft assemblies are poorly suited for this purpose because they miss repetitive sequences or genes that are central to the vector’s biology and evolution. Incomplete information about the correct copies or sequence of a gene may mislead conclusions about functional significance of the gene or the target sequence [15] and may lead to mistargeting or misuse in a gene drive. The *An. stephensi* reference assembly solves these problems, revealing previously invisible or uncharacterized structural and functional genomic elements that shape various aspects of the vector biology of *An. stephensi*. Additionally, functionally-important SVs are segregating even in this inbred lab strock, indicating a significant role of structural genetic variation in phenotypic variation in this species. Finally, recent advances in technology have sparked enthusiasm for sequencing all eukaryotes in the tree of life [25]. The assembly we report here is timely, as it constitutes the first malaria vector to reach the exacting reference standards called for by these ambitious proposals, and will stand alongside established references like those for human and fruit flies (fig. S5). This new assembly of *An. stephensi* provides a comprehensive and accurate map of genomic functional elements and will serve as a foundation for the new age of active genetics in *An. stephensi*.

## Materials and methods

### Mosquitoes

*Anopheles stephensi* mosquitoes of a strain (UCISS2018) from the Indian subcontinent (gift of M. Jacobs-Lorena, Johns Hopkins University)[68] were maintained in insectary conditions (27°C and 77% humidity) with a photoperiod of 12 h light:12h dark including 30 minutes of dawn and dusk at the University of California, Irvine (UCI). Larvae were reared in distilled water and fed ground TetraMin^®^ fish food mixed with yeast powder. Adults had unlimited access to sucrose solutions (10% wt/vol) and females were provided with blood meals consisting of defibrinated calf blood (Colorado Serum Co., Denver) through the Hemotek^®^ membrane feeding system. We established an isofemale line from the colony and inbred the line by sib mating for 5 generations prior to sequencing.

### Genome sequencing

Genomic DNA from 70 adult male and female mosquitoes was extracted using the Qiagen Blood & Cell Culture DNA Midi Kit following the previously-described protocol [69]. The genomic DNA was sheared with 10 plunges of size 21 blunt needles, followed by 10 plunges of size 24 blunt end needles. We generated our PacBio reads using 31 SMRTcells on the RSII platform (P6-C4 chemistry) at the UC San Diego Genomics Core and 2 SMRTcells on Sequel I platform at Nucleome (Hyderabad, India). From the same genomic DNA, we also generated 3.7 GB of 300 bp paired-end reads at the UC San Diego genomics core and 32.37 GB of 150 bp paired-end Illumina reads from Nucleome. To identify the Y-linked contigs, we generated 27.8 GB and 28.5 GB 100 bp paired-end Illumina reads from male and female genomic DNA, respectively, at UCI Genomics High-Throughput Facility (GHTF).

### RNA extraction and sequencing

Total RNA was extracted from a total of six samples prepared from pooled individuals: 5-7-day-old, sugar-fed males, 5-7-day-old, sugar-fed females, and blood-fed females 3-6 h, 24 h, 48 h, and 72 h after feeding. The male pool consisted of 15 individuals, while female pools comprised 10 individuals each. All samples were isolated from the same mosquito cage. To do so, sugar-fed male and female samples were collected, and a blood meal offered for 1 h. Unfed females were removed from the cage and blood-fed females retrieved at each of the indicated time points. At the time of collection, samples were immersed in 500 μL of RNAlater RNA Stabilization Reagent (Qiagen) and stored at 4°C. Total RNA was extracted using the RNeasy Mini Kit (Qiagen) following manufacturer’s instructions for the Purification of Total RNA from Animal Tissues. Extracted samples were treated with DNA-free Kit (Ambion) to remove traces of genomic DNA. Finally, samples were cleaned using the RNA Clean & Concentrator Kit (Zymo Research). mRNA selection, cDNA synthesis and Iso-Seq library prep was performed at UCI GHTF following the manufacturer’s (Pacific Biosciences) protocol. For each of the six samples, one SMRTcell of Iso-seq reads were generated on the Sequel I platform.

### Genome assembly

We used 42.4 GB or 180× of long reads (assuming haploid genome size G = 235 Mb) to generate two draft assemblies of *An. stephensi* using Canu v1.7 [70] and Falcon v2.1.4 [71]. Falcon was used to assemble the heterozygous regions (fig. S1) that were recalcitrant to Canu. We filled the gaps in the canu assembly using the Falcon primary contigs following the two-steps merging approach with Quickmerge v0.3, where the Canu assembly was used as the reference assembly in the first merging step [69, 72]. The resulting assembly was processed with finisherSC (v2.1) to remove the redundant contigs and to fill the further gaps with raw reads [73]. This PacBio assembly (613 contigs, contig N50 = 38.1 Mb, 257.1 Mb in total) was polished twice with arrow (smrtanalysis v5.2.1) and twice with Pilon v1.22 using ~400X (80 Gb) 150 bp PE Illumina reads from our three Illumina dataset [74].

### Identification of polymorphic mutations

To identify the variants segregating in the sequenced strain, we aligned the alternate haplotype contigs (a_ctg.fa) identified by Falcon to the scaffolded assembly. Then we called the indels using SVMU v0.2 (Structural Variants from MUmmer). An indel was marked as a TE based on its overlap with the Repeatmasker annotated TEs. To estimate heterozygosity, we mapped the Illumina reads to the chromosome scaffolds using bowtie2 (v2.2.7) and converted the alignments to a sorted bam file using SAMtools (v1.9). A VCF file containing the SNPs and small indels were generated using freebayes (v1.3.2-40-gcce27fc) and pairwise nucleotide diversity (pi) was calculated over 25 kb windows using vcftools (vcftools --window-pi 25000; v0.1.14v0.1.14).

### Microbial sequence decontamination

Microbial contigs in the assembly were identified using Kraken v2.0.7-beta [75], which assigned taxonomic labels to the 613 contigs (fig. S4). Kraken mapped k-mers (31-35 nt default) from the 613 contig sequences against the databases from the six domain sets: bacteria, archaea, viral, UniVec_Core, fungi, and protozoa from National Center for Biotechnology Information (NCBI) and the genome sets including representative reference mosquito from VectorBase v2019-02 and *Drosophila* genomes (n=24; table S9). The databases map k-mers to the lowest common ancestor (LCA) of all genomes known to contain a given k-mer. Kraken label for each contig was further classified as either *Anopheles*, contaminating (non-*Anopheles*), or unclassified (no hit in the database) (fig. S4). To prevent false positives in the results, low-complexity sequences in the assembly were masked with dustmasker (blast v2.8.1) [76] prior to running Kraken. The mitochondrial genome of *An. stephensi* was identified by aligning the existing mitogenome (GenBank No. KT899888) against the contigs using nucmer in MUMmer v4.0.0b [77].

### Scaffolding

To *de novo* scaffold the microbial decontaminated 566 contigs, we collected HiC data from adult male and female mosquitoes filled up to the 1ml mark of an 1.5 ml Eppendorf tube. We flash-froze the adult mosquitoes and sent them to Arima Genomics (San Diego) to generate a HiC library using the Arima kit. This library was sequenced on a single flow cell of an Illumina HiSeq 2500 instrument, generating 326 GB of Illumina 150 bp paired-end reads. We mapped the HiC reads to the *An. stephensi* contigs using Juicer v1.5.6 [78] and used the resulting contact map to scaffold the contigs using 3D-DNA v180922 [79]. The order and orientation of the three chromosomes were examined by nucmer in MUMmer v4.0.0b alignment of 20 gene/probe physical map data (X, 5 probes; 2, 7; 3, 8) generated from Fluorescence In Situ Hybridization (FISH) on polytene chromosomes (fig. S10; table S10) [18] against Hi-C chromosome assemblies.

### QV estimation and assembly statistics

To estimate the error rate in our final assembly, we mapped the paired-end Illumina reads to the assembly using bwa mem (bwa v0.7.17-5). The alignments were converted to bam format and then sorted using SAMtools (v.1.8-11). We called the variants using freeBayes v0.9.21 [80] (command: freebayes -C 2 -0 -O -q 20 -z 0.10 -E 0 -X -u -p 2 -F 0.75) and followed the approach of [70] to calculate QV (10*-log10(2981/250,632,892)). Briefly, we counted the number of bases comprising homozygous variants in the assembly (2981) and then divided it by the total mapped bases that had a coverage of at least three (250,632,892). We used QUAST v5.0.3 to obtain assembly statistics [81]. Out of 250 Mb, 205 Mb (82%) were scaffolded into the three chromosome-length scaffolds that correspond to the three *An. stephensi* chromosomes (chrX, 22.7 Mb; chr2, 93.7 Mb; chr3, 88.7 Mb). We identified 66 unplaced contigs as alternate haplotigs (66 contigs = 7.2 Mb) using mummer alignments of contigs to the major chromosomes. Additionally, 103 (8.6 Mb) of 458 (35.9 Mb) unplaced or unclassified contigs were identified as alternate haplotigs using BUSCO v4.1.4 Diptera odb10 dataset [82] and the software Purge_dups v1.0 [83] (Table 1) (table S12). The final Hi-C map was visualized using HiCExplorer v3.4.2 [84]. We estimated the proportion of scaffolded haploid genomes using the publicly available resources on *An. stephensi* and *An. gambiae* genome size [85, 86]. The C-value of *An. stephensi* is 0.24 and that of *An. gambiae* is 0.27. Based on the odds ratio of the genome sizes of the two species estimated from their C-values and the *An. gambiae* genome size (265 Mb), we inferred the genome size for *An. stephensi* to be ~235 Mb.

### Repeat annotation

We created a custom TE library using the EDTA (Extensive *de-novo* TE Annotator) pipeline [87] and Repeatmodeler v2.0.0 (http://www.repeatmasker.org/RepeatModeler/) to annotate the TEs. LTR retrotransposons and DNA elements were identified *de novo* using EDTA, but because EDTA does not identify Non-LTR elements, Repeatmodeler was used to identify these. The two libraries were combined and the final library was used with Repeatmasker (v4.0.7) to annotate the genome-wide TEs. Tandem repeats were annotated using Tandem Repeat Finder v4.09 [88]. The number and copy number of micro-, mini- and macro-satellites spanning in each 100 kb non-overlapping window of the three chromosomes were identified. The satellite classification was made as described in [18]. In brief, tandem repeats were classified as micro-(1-6 bases), mini-(7-99 bases) and macro-(>=100 bases) satellites. Mini- and macro-satellites were considered only if they had a copy number of more than 2. All these three simple repeats were considered only if they had at least 80% sequence identity, and set some cutoff (>=2 copy number; >=80% identity) to screen high confidence repeats, then the overall abundance was calculated.

### Annotation using Iso-Seq

In total, six samples (5 females; 1 male) of *An. stephensi* mosquitoes were used for Iso-Seq sequencing (see RNA extraction and sequencing). Raw PacBio long-molecule sequencing data was processed using the SMRT analysis v7.0.0 Iso-Seq3 pipeline [89] (supplemental Text). Briefly, CCS was used to generate the full-length (FL) reads for which all 5’-end primer, polyA tail and 3’-end primer have been sequenced and then Lima was used to identify and remove the 5’- and 3’-end cDNA primers from the FL reads. The resulting bam files were processed with Iso-Seq3 to refine and cluster the reads, which were polished with Arrow. This *de novo* pipeline outputs FASTQ files containing two sets of error-corrected, full length isoforms: i) the high-quality set contains isoforms supported by at least two FL reads with an accuracy of at least 99% and ii) while the low-quality set contains isoforms with an accuracy <99% that occurred due to insufficient coverage or rare transcripts. The high quality isoforms were collapsed with Cupcake v10.0.1 and were used in Talon v5.0 for annotation [90]. We combined the high quality isoforms with other lines of evidence using MAKER2 v2.31.10 to create a final annotation (see below).

### MAKER Annotation

The final annotation of the genome was performed using MAKER2 v2.31.10 [91], which combines empirical evidence and *ab initio* gene prediction to produce final annotations (supplemental Text). We used MAKER2 for three cycles of gene predictions. First, the Iso-Seq data were used as evidence for training MAKER2 for gene predictions. We also used transcriptome and peptide sequence data from *An. gambiae* (PEST4.12) and *An. funestus* (FUMOZ 3.1) as alternative evidence to support the predicted gene models. Prior to gene annotation, repeats were masked using RepeatMasker included in MAKER2. Mapping of EST and protein evidence to the genome by MAKER2 using BLASTn and BLASTx, respectively, yielded 12,324 genes transcribing 14,888 mRNAs.

The output of first round gene models were used for the second round, where MAKER2 ran SNAP and AUGUSTUS for *ab initio* gene predictions. Next, another round of SNAP and AUGUSTUS predictions were performed to synthesize the final annotations that produced 14,966 genes, transcribing 16,559 mRNAs. In total, we identified 56,388 exons, 9,791 5’-end UTRs, 9290 3’end UTRs and 503 tRNAs (Table 1; see Methods). We also predicted *ab initio* an additional 14,192 mRNAs/proteins but due to weak support they were not considered. The final MAKER annotation was assessed using recommended AED and Pfam statistical metrics.

The gene models were functionally annotated in MAKER2 v2.31.10 through a homology BLAST search to UniProt-Sprot database, while the domains in the annotated proteins were assigned from the InterProScan database (supplementary Text). We compared our gene model annotations with the draft assembly using OrthoFinder v2.3.7 [92] The GO enrichment analysis was performed in PANTHER v15.0 using PANTHER GO-SLIM Biological Process annotation data set [93]. Further, the orthologous top 1% of *An. stephensi* up-regulated gene protein sequences in *An. gambiae* were identified by OrthoFinder v2.3.7. The final MAKER2 GFF file, the list of insecticide resistance genes, the list of immunity genes, and the list of novel transcripts are available at https://github.com/mahulchak/stephensi_genome.

### Validation and quantification with RNAseq and Iso-Seq

To quantify transcript abundance using Iso-seq reads, raw reads were mapped to the genome assembly using minimap2 [94] and the gene-specific transcript abundance was measured using bedtools, requiring that each Iso-seq read overlaps at least 75% of gene length annotated with TALON v5.0 (bedtools coverage -mean -F 0.75 -a talon.gff-b minimap.bam) [95]. To take variation due to sequencing yield per SMRTcell into account while calculating transcript abundance, Iso-seq coverage of each gene was divided by a normalization factor, which was calculated by dividing the total read counts for each sample by the total read counts from the unfed female sample. To identify the genes up- or down-regulated due to blood feeding, we compared the transcript abundance of 5-7 days old adult females before and after blood meal. To identify the genes whose expressions are most strongly affected by blood meal, we rank-ordered transcript abundance differences for all genes showing non-zero transcript abundance in either of the two samples. Here we reported the genes that are in the top 1% of the transcript level differences which corresponds to all genes showing >~64-fold increase or decrease in transcript abundance between the two samples.

To obtain transcript levels of Y-linked genes from embryos, larvae, and adults, we used publicly available RNA-seq data (table S11). RNA-seq reads were mapped to the genome using HISAT2 and the per-base read coverage was calculated from the sorted bam files using samtools depth. Additionally, the bam files were processed with stringtie to generate sample-specific transcript annotation in GTF format [96]. Sample specific GTF files were merged with stringtie to generate the final GTF. To obtain the gene model and transcript isoforms of *kdr*, stringtie annotated transcripts that covered the entire predicted ORF based on homology with *D. melanogaster para* were used.

### Identification of new genes and repeat elements

To identify incomplete or absent sequences in the most contiguous *An. stephensi* published assembly [18], the contigs from the draft assembly were aligned to the new assembly using nucmer [77] and alignments due to repeats were filtered using delta-filter to generate 1-to-1 mapping (delta-filter -r -q) between the two assemblies. The resulting delta file was converted into tab-separated alignment format using show-cords utility in MUMmer (v4). To identify TE sequences that are present in our assembly but absent in Jiang et al. (2014), we annotated the TEs in the latter using RepeatMasker and calculated the abundance of different classes of TEs (DNA, LTR, Long Interspersed Nuclear Elements or Non-LTR) from the RepeatMasker output. Additionally, we identified TE sequences in our assembly that were either fragmented or absent in the Jiang et al. (2014) contig assembly by looking for TE sequences that either failed to map or mapped only partially to the latter (bedtools intersect -v -f 1.0 -a te.bed -b alignment.bed). To identify the genes that were fragmented or absent from the Jiang et al. [18] data, we combined two complementary approaches. First, we used the 1-to-1 genome alignment to identify genes that were split over >1 contigs. Second, we mapped the annotated protein sequences from our study to the protein sequences reported in Jiang et al. using OrthoFinder v.2.3.7 [92] and identified the transcripts that were present only in our study. We combined the unique genes found by each approach to calculate the total number of previously incomplete or missing genes. To rule out assembly errors in the new assembly as the cause of discrepancy between the two assemblies, 20 features disagreeing between the assemblies were randomly selected from each category and manually inspected in IGV. At least 3 long reads spanning an entire feature was used as evidence for correct assembly of the features.

### Identification of Y contigs

To identify the putative Y contigs, male and female specific k-mers were identified from male and female paired-end Illumina reads using Jellyfish [97] (fig. S11). Density of male and female k-mers for each contig was calculated and the contigs showing more than two-fold higher density of male specific k-mers were designated as putative Y contigs. Interestingly, the *Serratia* genome we assembled also showed similar male k-mer enrichment as the Y contigs.

### Experimental validation of Y-linked contigs

The k-mer based approach employed to identify male-specific kmers that occur at a rate 20-fold higher than female-specific kmers in the *An. stephensi* scaffolds (fig. S11). In order to verify putative Y-linked sequences, ten 2-3 days-old male or female *An. stephensi* mosquitoes per replicate were used for the experiment. Genomic DNA was extracted from each sample using DNeasy Blood and Tissue Kit (Cat # 69504). Gene specific primers (Y15 forward(F) - ATT TTA GTT ATT TAG AGG CTT CGA, Y15 reverse(R) - GCG TAT GAT AGA AAC CGC AT; Y22 F - ATG CCA AAA AAA CGG TTG CG, Y22 R - CTA GCT CTT GTA AAG AGT CAC CTT; Y28 F - ATG CTA CAA AAC AGT GCC TT, Y28 R - TTA GGT CAG ATA TAG ACA CAG ACA CA) were designed based on the genome sequence to amplify ≥500 bp products using Polymerase Chain Reaction (PCR) reaction. The amplification was done using Q5 high fidelity 2X Master Mix (Cat # M0492). Amplicons were resolved in agarose gels and male versus female amplification was compared. The PCR products were gel eluted and Sanger sequenced (fig. S11) (Genewiz) with forward PCR primer. The identity of the sequencing was confirmed by aligning the amplicon sequences against the *An. stephensi* genome assembly using BLAST.

### Identification of putative immune gene families

Studying the patterns of evolution in innate immune genes facilitate understanding the evolutionary dynamics of *An. stephensi* and pathogens they harbor. A total of 1649 manually curated immune proteins of *An. gambiae* (Agam 385)*, Ae. aegypti* (Aaeg 422), *Cu. quinquefasciatus* (Cpip 495) and *D. melanogaster* (Dmel 347) in ImmunoDB (table S8) [98] were used as databases to search for the putative immune-related proteins in MAKER2-annotated protein sequences of the *An. stephensi* assembly using sequence alignment and phylogenetic orthology inference based method in OrthoFinder v2.3.7. The number of single copy orthogroup/orthologous proteins (one-to-one) and co-orthologous and paralogous proteins in *An. stephensi* were identified (one-to-many; many-to-one; many-to-many).

## Declarations

### Ethics approval and consent to participate

Not applicable

### Consent for publication

Not applicable

### Availability of data and materials

The sequenced strain is available at no cost from AJ. The raw PacBio, illumina and Hi-C sequencing data and *An. stephensi* genome assembly were deposited in the NCBI BioProject database (Accession number PRJNA629843). The annotations and other genomic features can be accessed at http://3.93.125.130/tigs/anstephdb/. All codes used in the study, including those used to make figures are available at https://github.com/mahulchak/stephensi_genome.

### Authors’ contributions

MC, AAJ, and JJE conceived the experimental approach; AA, PH, BK and LTN performed laboratory research; MC, AR, SJ, KP, and SW performed bioinformatics analysis; SSr coordinated all activities at IBAB and contributed to the annotation pipeline. MC and AR wrote the manuscript draft; EB, SSu, AAJ and JJE edited the draft. SSu coordinated the project activities between TIGS-India and TIGS-UC San Diego, and was involved in planning of the sequencing strategies. All authors contributed to the finalized version of the manuscript.

## Acknowledgements

We thank Judith Coleman for help with mosquito collection and Yi Liao for helpful discussions.

## References

1. Roberts L. Mosquitoes and disease. Science. 2002;298:82–3.

2. Institute of Medicine (US) Committee for the Study on Malaria Prevention and Control, Oaks SC Jr, Mitchell VS, Pearson GW, Carpenter CCJ. Vector Biology, Ecology, and Control. National Academies Press (US); 1991.

3. Cohuet A, Harris C, Robert V, Fontenille D. Evolutionary forces on Anopheles: what makes a malaria vector? Trends Parasitol. 2010;26:130–6.

4. Holt RA, Subramanian GM, Halpern A, Sutton GG, Charlab R, Nusskern DR, et al. The genome sequence of the malaria mosquito Anopheles gambiae. Science. 2002;298:129–49.

5. Severson DW, Behura SK. Mosquito genomics: progress and challenges. Annu Rev Entomol. 2012;57:143–66.

6. The Anopheles gambiae 1000 Genomes Consortium, Clarkson CS, Miles A, Harding NJ, Lucas ER, Battey CJ, et al. Genome variation and population structure among 1,142 mosquitoes of the African malaria vector species Anopheles gambiae and Anopheles coluzzii. bioRxiv. 2019;:864314. doi:10.1101/864314.

7. Neafsey DE, Waterhouse RM, Abai MR, Aganezov SS, Alekseyev MA, Allen JE, et al. Mosquito genomics. Highly evolvable malaria vectors: the genomes of 16 Anopheles mosquitoes. Science. 2015;347:1258522.

8. Waterhouse RM, Aganezov S, Anselmetti Y, Lee J, Ruzzante L, Reijnders MJMF, et al. Evolutionary superscaffolding and chromosome anchoring to improve Anopheles genome assemblies. BMC Biol. 2020;18:1.

9. Sharma VP. Current scenario of malaria in India. Parassitologia. 1999;41:349–53.

10. Seyfarth M, Khaireh BA, Abdi AA, Bouh SM, Faulde MK. Five years following first detection of Anopheles stephensi (Diptera: Culicidae) in Djibouti, Horn of Africa: populations established-malaria emerging. Parasitol Res. 2019;118:725–32.

11. Sinka ME, Pironon S, Massey NC, Longbottom J, Hemingway J, Moyes CL, et al. A new malaria vector in Africa: Predicting the expansion range of Anopheles stephensi and identifying the urban populations at risk. Proceedings of the National Academy of Sciences. 2020;:202003976. doi:10.1073/pnas.2003976117.

12. James AA. Gene drive systems in mosquitoes: rules of the road. Trends Parasitol.2005;21:64–7.

13. Gantz VM, Jasinskiene N, Tatarenkova O, Fazekas A, Macias VM, Bier E, et al. Highly efficient Cas9-mediated gene drive for population modification of the malaria vector mosquito Anopheles stephensi. Proc Natl Acad Sci U S A. 2015;112:E6736–43.

14. Unckless RL, Clark AG, Messer PW. Evolution of Resistance Against CRISPR/Cas9 Gene Drive. Genetics. 2017;205:827–41.

15. Chakraborty M, VanKuren NW, Zhao R, Zhang X, Kalsow S, Emerson JJ. Hidden genetic variation shapes the structure of functional elements in Drosophila. Nat Genet. 2018;50:20–5.

16. Treangen TJ, Salzberg SL. Repetitive DNA and next-generation sequencing: computational challenges and solutions. Nat Rev Genet. 2011;13:36–46.

17. Catteruccia F, Nolan T, Loukeris TG, Blass C, Savakis C, Kafatos FC, et al. Stable germline transformation of the malaria mosquito Anopheles stephensi. Nature. 2000;405:959–62.

18. Jiang X, Peery A, Hall AB, Sharma A, Chen X-G, Waterhouse RM, et al. Genome analysis of a major urban malaria vector mosquito, Anopheles stephensi. Genome Biol. 2014;15:459.

19. Matthews BJ, Dudchenko O, Kingan SB, Koren S, Antoshechkin I, Crawford JE, et al. Improved reference genome of Aedes aegypti informs arbovirus vector control. Nature. 2018;563:501–7.

20. Hoskins RA, Carlson JW, Wan KH, Park S, Mendez I, Galle SE, et al. The Release 6 reference sequence of the Drosophila melanogaster genome. Genome Res. 2015;25:445–58.

21. Ghurye J, Koren S, Small ST, Redmond S, Howell P, Phillippy AM, et al. A chromosome-scale assembly of the major African malaria vector Anopheles funestus. GigaScience. 2019;8. doi:10.1093/gigascience/giz063.

22. Kingan SB, Heaton H, Cudini J, Lambert CC, Baybayan P, Galvin BD, et al. A High-Quality De novo Genome Assembly from a Single Mosquito Using PacBio Sequencing. Genes. 2019;10. doi:10.3390/genes10010062.

23. Compton A, Liang J, Chen C, Lukyanchikova V, Qi Y, Potters M, et al. The Beginning of the End: A Chromosomal Assembly of the New World Malaria Mosquito Ends with a Novel Telomere. G3. 2020;10:3811–9.

24. Lukyanchikova V, Nuriddinov M, Belokopytova P, Liang J, Reijnders MJMF, Ruzzante L, et al. Anopheles mosquitoes revealed new principles of 3D genome organization in insects. Genomics. 2020;:4912.

25. Lewin HA, Robinson GE, Kress WJ, Baker WJ, Coddington J, Crandall KA, et al. Earth BioGenome Project: Sequencing life for the future of life. Proc Natl Acad Sci U S A. 2018;115:4325–33.

26. Hall AB, Papathanos P-A, Sharma A, Cheng C, Akbari OS, Assour L, et al. Radical remodeling of the Y chromosome in a recent radiation of malaria mosquitoes. Proc Natl Acad Sci U S A. 2016;113:E2114–23.

27. Wang S, Dos-Santos ALA, Huang W, Liu KC, Oshaghi MA, Wei G, et al. Driving mosquito refractoriness to Plasmodium falciparum with engineered symbiotic bacteria. Science. 2017;357:1399–402.

28. Ribeiro JM, Kidwell MG. Transposable elements as population drive mechanisms: specification of critical parameter values. J Med Entomol. 1994;31:10–6.

29. Macias VM, Jimenez AJ, Burini-Kojin B, Pledger D, Jasinskiene N, Phong CH, et al. nanos-Driven expression of piggyBac transposase induces mobilization of a synthetic autonomous transposon in the malaria vector mosquito, Anopheles stephensi. Insect Biochem Mol Biol. 2017;87:81–9.

30. Jasinskiene N, Coates CJ, Benedict MQ, Cornel AJ, Rafferty CS, James AA, et al. Stable transformation of the yellow fever mosquito, Aedes aegypti, with the Hermes element from the housefly. Proc Natl Acad Sci U S A. 1998;95:3743–7.

31. Arensburger P, Kim Y-J, Orsetti J, Aluvihare C, O’Brochta DA, Atkinson PW. An active transposable element, Herves, from the African malaria mosquito Anopheles gambiae. Genetics. 2005;169:697–708.

32. Chakraborty M, Chang C-H, Khost DE, Vedanayagam J, Adrion JR, Liao Y, et al. Evolution of genome structure in the Drosophila simulans species complex. bioRxiv. 2020;:2020.02.27.968743. doi:10.1101/2020.02.27.968743.

33. Volkenhoff A, Weiler A, Letzel M, Stehling M, Klämbt C, Schirmeier S. Glial Glycolysis Is Essential for Neuronal Survival in Drosophila. Cell Metab. 2015;22:437–47.

34. Zhou S, Luoma SE, St Armour GE, Thakkar E, Mackay TFC, Anholt RRH. A Drosophila model for toxicogenomics: Genetic variation in susceptibility to heavy metal exposure. PLoS Genet. 2017;13:e1006907.

35. Robertson SE, Dockendorff TC, Leatherman JL, Faulkner DL, Jongens TA. germ cell-less is required only during the establishment of the germ cell lineage of Drosophila and has activities which are dependent and independent of its localization to the nuclear envelope. Dev Biol. 1999;215:288–97.

36. Prowse TA, Adikusuma F, Cassey P, Thomas P, Ross JV. A Y-chromosome shredding gene drive for controlling pest vertebrate populations. Elife. 2019;8. doi:10.7554/eLife.41873.

37. Muller HJ. THE RELATION OF RECOMBINATION TO MUTATIONAL ADVANCE. Mutat Res. 1964;106:2–9.

38. Hall AB, Qi Y, Timoshevskiy V, Sharakhova MV, Sharakhov IV, Tu Z. Six novel Y chromosome genes in Anopheles mosquitoes discovered by independently sequencing males and females. BMC Genomics. 2013;14:273.

39. Kokoza V, Ahmed A, Woon Shin S, Okafor N, Zou Z, Raikhel AS. Blocking of Plasmodium transmission by cooperative action of Cecropin A and Defensin A in transgenic Aedes aegypti mosquitoes. Proc Natl Acad Sci U S A. 2010;107:8111–6.

40. Shane JL, Grogan CL, Cwalina C, Lampe DJ. Blood meal-induced inhibition of vector-borne disease by transgenic microbiota. Nat Commun. 2018;9:4127.

41. Muller HM, Catteruccia F, Vizioli J, Dellatorre A, Crisanti A. Constitutive and Blood Meal-Induced Trypsin Genes in Anopheles gambiae. Exp Parasitol. 1995;81:371–85.

42. Marinotti O, Nguyen QK, Calvo E, James AA, Ribeiro JMC. Microarray analysis of genes showing variable expression following a blood meal in Anopheles gambiae. Insect Mol Biol. 2005;14:365–73.

43. Hammond A, Galizi R, Kyrou K, Simoni A, Siniscalchi C, Katsanos D, et al. A CRISPR-Cas9 gene drive system targeting female reproduction in the malaria mosquito vector Anopheles gambiae. Nat Biotechnol. 2016;34:78–83.

44. Rahman MM, Franch-Marro X, Maestro JL, Martin D, Casali A. Local Juvenile Hormone activity regulates gut homeostasis and tumor growth in adult Drosophila. Sci Rep. 2017;7:11677.

45. Taracena ML, Bottino-Rojas V, Talyuli OAC, Walter-Nuno AB, Oliveira JHM, Angleró-Rodriguez YI, et al. Regulation of midgut cell proliferation impacts Aedes aegypti susceptibility to dengue virus. PLoS Negl Trop Dis. 2018;12:e0006498.

46. Vizioli J, Bulet P, Charlet M, Lowenberger C, Blass C, Müller HM, et al. Cloning and analysis of a cecropin gene from the malaria vector mosquito, Anopheles gambiae. Insect Mol Biol. 2000;9:75–84.

47. Vatandoost H, Hanafi-Bojd AA. Indication of pyrethroid resistance in the main malaria vector, Anopheles stephensi from Iran. Asian Pac J Trop Med. 2012;5:722–6.

48. Safi NHZ, Ahmadi AA, Nahzat S, Warusavithana S, Safi N, Valadan R, et al. Status of insecticide resistance and its biochemical and molecular mechanisms in Anopheles stephensi (Diptera: Culicidae) from Afghanistan. Malar J. 2019;18:249.

49. Yared S, Gebressielasie A, Damodaran L, Bonnell V, Lopez K, Janies D, et al. Insecticide Resistance in Anopheles stephensi in Somali Region, Eastern Ethiopia. In Review. 2019.

50. Enayati AA, Vatandoost H, Ladonni H, Townson H, Hemingway J. Molecular evidence for a kdr-like pyrethroid resistance mechanism in the malaria vector mosquito Anopheles stephensi. Med Vet Entomol. 2003;17:138–44.

51. Pedra JHF, McIntyre LM, Scharf ME, Pittendrigh BR. Genome-wide transcription profile of field- and laboratory-selected dichlorodiphenyltrichloroethane (DDT)-resistant Drosophila. Proc Natl Acad Sci U S A. 2004;101:7034–9.

52. Mouchès C, Pasteur N, Bergé JB, Hyrien O, Raymond M, de Saint Vincent BR, et al. Amplification of an esterase gene is responsible for insecticide resistance in a California Culex mosquito. Science. 1986;233:778–80.

53. Cridland JM, Macdonald SJ, Long AD, Thornton KR. Abundance and Distribution of Transposable Elements in Two Drosophila QTL Mapping Resources. Mol Biol Evol. 2013;30:2311–27.

54. Chakraborty M, Emerson JJ, Macdonald SJ, Long AD. Structural variants exhibit widespread allelic heterogeneity and shape variation in complex traits. Nat Commun. 2019;10:4872.

55. Qi Y, Wu Y, Saunders R, Chen X-G, Mao C, Biedler JK, et al. Guy1, a Y-linked embryonic signal, regulates dosage compensation in Anopheles stephensi by increasing X gene expression. Elife. 2019;8. doi:10.7554/eLife.43570.

56. Bhaskar V, Courey AJ. The MADF-BESS domain factor Dip3 potentiates synergistic activation by Dorsal and Twist. Gene. 2002;299:173–84.

57. Fakhouri M, Elalayli M, Sherling D, Hall JD, Miller E, Sun X, et al. Minor proteins and enzymes of the Drosophila eggshell matrix. Dev Biol. 2006;293:127–41.

58. Amenya DA, Chou W, Li J, Yan G, Gershon PD, James AA, et al. Proteomics reveals novel components of the Anopheles gambiae eggshell. J Insect Physiol. 2010;56:1414–9.

59. Ahmed SMH, Maldera JA, Krunic D, Paiva-Silva GO, Pénalva C, Teleman AA, et al. Fitness trade-offs incurred by ovary-to-gut steroid signalling in Drosophila. Nature. 2020. doi:10.1038/s41586-020-2462-y.

60. Gayathri V, Murthy PB. Reduced susceptibility to deltamethrin and kdr mutation in Anopheles stephensi Liston, a malaria vector in India. J Am Mosq Control Assoc. 2006;22:678–88.

61. Prasad KM, Raghavendra K, Verma V, Velamuri PS, Pande V. Esterases are responsible for malathion resistance in Anopheles stephensi: A proof using biochemical and insecticide inhibition studies. J Vector Borne Dis. 2017;54:226–32.

62. Reimer L, Fondjo E, Patchoké S, Diallo B, Lee Y, Ng A, et al. Relationship Between kdr Mutation and Resistance to Pyrethroid and DDT Insecticides in Natural Populations of Anopheles gambiae. J Med Entomol. 2014;45:260–6.

63. Dykes CL, Kushwah RBS, Das MK, Sharma SN, Bhatt RM, Veer V, et al. Knockdown resistance (kdr) mutations in Indian Anopheles culicifacies populations. Parasites & Vectors. 2015;8. doi:10.1186/s13071-015-0946-7.

64. Hollister JD, Gaut BS. Epigenetic silencing of transposable elements: a trade-off between reduced transposition and deleterious effects on neighboring gene expression. Genome Res. 2009;19:1419–28.

65. Hemingway J, Hawkes NJ, McCarroll L, Ranson H. The molecular basis of insecticide resistance in mosquitoes. Insect Biochem Mol Biol. 2004;34:653–65.

66. Gantz VM, Bier E. The dawn of active genetics. Bioessays. 2016;38:50–63.

67. Carballar-Lejarazú R, James AA. Population modification of Anopheline species to control malaria transmission. Pathog Glob Health. 2017;111:424–35.

68. Nirmala X, Marinotti O, Sandoval JM, Phin S, Gakhar S, Jasinskiene N, et al. Functional characterization of the promoter of the vitellogenin gene, AsVg1, of the malaria vector, Anopheles stephensi. Insect Biochem Mol Biol. 2006;36:694–700.

69. Chakraborty M, Baldwin-Brown JG, Long AD, Emerson JJ. Contiguous and accurate de novo assembly of metazoan genomes with modest long read coverage. Nucleic Acids Res. 2016;44:e147.

70. Koren S, Walenz BP, Berlin K, Miller JR, Bergman NH, Phillippy AM. Canu: scalable and accurate long-read assembly via adaptive k-mer weighting and repeat separation. Genome Res. 2017. doi:10.1101/gr.215087.116.

71. Chin C-S, Alexander DH, Marks P, Klammer AA, Drake J, Heiner C, et al. Nonhybrid, finished microbial genome assemblies from long-read SMRT sequencing data. Nat Methods. 2013;10:563–9.

72. Solares EA, Chakraborty M, Miller DE, Kalsow S, Hall K, Perera AG, et al. Rapid Low-Cost Assembly of the Drosophila melanogaster Reference Genome Using Low-Coverage, Long-Read Sequencing. G3. 2018. doi:10.1534/g3.118.200162.

73. Lam K-K, LaButti K, Khalak A, Tse D. FinisherSC: a repeat-aware tool for upgrading de novo assembly using long reads. Bioinformatics. 2015;31:3207–9.

74. Walker BJ, Abeel T, Shea T, Priest M, Abouelliel A, Sakthikumar S, et al. Pilon: An Integrated Tool for Comprehensive Microbial Variant Detection and Genome Assembly Improvement. PLoS One. 2014;9:e112963.

75. Wood DE, Salzberg SL. Kraken: ultrafast metagenomic sequence classification using exact alignments. Genome Biol. 2014;15:R46.

76. Morgulis A, Coulouris G, Raytselis Y, Madden TL, Agarwala R, Schäffer AA. Database indexing for production MegaBLAST searches. Bioinformatics. 2008;24:1757–64.

77. Marçais G, Delcher AL, Phillippy AM, Coston R, Salzberg SL, Zimin A. MUMmer4: A fast and versatile genome alignment system. PLoS Comput Biol. 2018;14:e1005944.

78. Durand NC, Shamim MS, Machol I, Rao SSP, Huntley MH, Lander ES, et al. Juicer Provides a One-Click System for Analyzing Loop-Resolution Hi-C Experiments. Cell Syst. 2016;3:95–8.

79. Dudchenko O, Batra SS, Omer AD, Nyquist SK, Hoeger M, Durand NC, et al. De novo assembly of the Aedes aegypti genome using Hi-C yields chromosome-length scaffolds. Science. 2017;356:92–5.

80. Garrison E, Marth G. Haplotype-based variant detection from short-read sequencing. arXiv [q-bio.GN]. 2012. http://arxiv.org/abs/1207.3907.

81. Mikheenko A, Saveliev V, Gurevich A. MetaQUAST: evaluation of metagenome assemblies. Bioinformatics. 2016;32:1088–90.

82. Waterhouse RM, Seppey M, Simão FA, Manni M, Ioannidis P, Klioutchnikov G, et al. BUSCO applications from quality assessments to gene prediction and phylogenomics. Mol Biol Evol. 2017. doi:10.1093/molbev/msx319.

83. Guan D, McCarthy SA, Wood J, Howe K, Wang Y, Durbin R. Identifying and removing haplotypic duplication in primary genome assemblies. Bioinformatics. 2020;36:2896–8.

84. Ramírez F, Bhardwaj V, Arrigoni L, Lam KC, Grüning BA, Villaveces J, et al. High-resolution TADs reveal DNA sequences underlying genome organization in flies. Nat Commun. 2018;9:189.

85. Jost E, Mameli M. DNA content in nine species of Nematocera with special reference to the sibling species of the Anopheles maculipennis group and the Culex pipiens group. Chromosoma. 1972;37:201–8.

86. Gregory TR. Animal Genome Size Database. http://www.genomesize.com. 2020.

87. Ou S, Su W, Liao Y, Chougule K, Agda JRA, Hellinga AJ, et al. Benchmarking transposable element annotation methods for creation of a streamlined, comprehensive pipeline. Genome Biol. 2019;20:275.

88. Benson G. Tandem repeats finder: a program to analyze DNA sequences. Nucleic Acids Res. 1999;27:573–80.

89. Gordon SP, Tseng E, Salamov A, Zhang J, Meng X, Zhao Z, et al. Widespread Polycistronic Transcripts in Fungi Revealed by Single-Molecule mRNA Sequencing. PLoS One. 2015;10:e0132628.

90. Wyman D, Balderrama-Gutierrez G, Reese F, Jiang S, Rahmanian S, Zeng W, et al. A technology-agnostic long-read analysis pipeline for transcriptome discovery and quantification. bioRxiv. 2019;:672931. doi:10.1101/672931.

91. Holt C, Yandell M. MAKER2: an annotation pipeline and genome-database management tool for second-generation genome projects. BMC Bioinformatics. 2011;12:491.

92. Emms DM, Kelly S. OrthoFinder: phylogenetic orthology inference for comparative genomics. Genome Biol. 2019;20:238.

93. Mi H, Muruganujan A, Ebert D, Huang X, Thomas PD. PANTHER version 14: more genomes, a new PANTHER GO-slim and improvements in enrichment analysis tools. Nucleic Acids Res. 2019;47:D419–26.

94. Li H. Minimap2: pairwise alignment for nucleotide sequences. Bioinformatics. 2018;34:3094–100.

95. Quinlan AR, Hall IM. BEDTools: a flexible suite of utilities for comparing genomic features. Bioinformatics. 2010;26:841–2.

96. Pertea M, Kim D, Pertea GM, Leek JT, Salzberg SL. Transcript-level expression analysis of RNA-seq experiments with HISAT, StringTie and Ballgown. Nat Protoc. 2016;11:1650–67.

97. Marçais G, Kingsford C. A fast, lock-free approach for efficient parallel counting of occurrences of k-mers. Bioinformatics. 2011;27:764–70.

98. Waterhouse RM, Kriventseva EV, Meister S, Xi Z, Alvarez KS, Bartholomay LC, et al. Evolutionary dynamics of immune-related genes and pathways in disease-vector mosquitoes. Science. 2007;316:1738–43.

