## Supplementary Information for "Hidden genomic features of an invasive malaria vector, *Anopheles stephensi*"

**This supplementary materials file includes:**

Supplementary text

figures S1 to S11

tables S1, S9, and S11

Legends for supplementary table files S2 to S8, S10 and S12

**Other supplementary materials for this manuscript include the following:**

Supplementary table files S2 to S8, S10 and S12

### Supplementary text

#### Microbial and mitochondrial sequences

*An. stephensi* genomic DNA was expected to contain microbial DNA [1] from endosymbionts [2], lab contaminations [3], and environment [4]. In total, 8% of the contigs (46/613) were from microbial sources and one contig (1/613) represented the mitochondrial genome (fig. S4). Complications from assembling a circular genome using softwares specializing on *de novo* assembly of linear genomes created three tandem copies of the mitochondrial genome in a 45 Kb contig. Manual trimming and curation produced a single complete mitogenome (15 Kb) that shared 99.8% sequence identity with a Chinese *An. stephensi* mitogenome (GenBank #KT899888.1). Among the 46 microbial contigs, 45 (1.3 Kb – 5.58 Mb) belonged to 12 bacterial species of three phyla: proteobacteria (37), bacteroidetes (7) and actinobacteria (1). Interestingly, the complete genomes of facultative bacteria *Serratia marcescens* (5.58 Mb in total; 3 contigs) and a double stranded DNA virus *Salmonella phage* (60.2 Kb, 1 contig) were assembled. These three *Serratia* contigs were scaffolded into a first circular complete genome of the facultative endosymbiont *Serratia marcescens* from *Anopheles* [5] (fig. S4). The members of the Proteobacteria *Serratia* and *Asaia* and Flavobacteria *Elizabethkingia* were also common among the microbial contigs. These are also found in *Anopheles gambiae* [6–8]. This *Salmonella phage* virus genome has GC content of 58%, which is genetically similar to the virus reported from sewage samples or sewage-contaminated river water samples from India (GenBank ID: KR296691) [9].

#### Assembly validation

The final genome assembly showed a total size of the assembly 250.63 Mb (N50, 88.7 Mb, GC content, 44.91%; L50, 2), in which, 205 Mb (82%) were scaffolded into the three chromosome-length scaffolds that correspond to the three *An. stephensi* chromosomes (chrX, 22.7 Mb; chr2, 93.7 Mb; chr3, 88.7 Mb) (Fig. 1). The order and orientation of the chromosomes were examined with 20 randomly chosen physical map data generated from FISH on polytene chromosomes [10]. The aligned probes showed 83-100% sequence identities to our assembly, confirming the chromosome nomenclature we used based on synteny (fig. S10; table S10). The polished final version of assembly is highly accurate, free of large mis-assemblies with estimates of error less than 1 per 83kb (*phred* QV score > 49.2).

#### Assessment of genome completeness by BUSCO

The BUSCO [11] based universal 3285 orthologs (Diptera database) analysis showed the completeness of the reference-quality *An. stephensi* genome (99.7%) that we assembled exceeds the previous assemblies (draft assembly 97.2%; Astel2 98.9%) (fig. S5). Comparison with the recent version of chromosome-level genome assemblies of other *Anopheles* species (range 97-99.6%), *Aedes aegypti* (97.6%), *Culex quinquefasciatus* (87.7%) and *Drosophila melanogaster* (99.5%) showed that the completeness of *An. stephensi* genome exceeds all these assemblies (fig. S5). Among the 3285 Diptera BUSCOs in *An. stephensi* assembly, 3276 (99.7%) were complete (3256 (99.2%) were single-copy; 18 (0.5%) were duplicated), 2 fragmented and 7 are missing BUSCOs. All complete BUSCOs were identified in major 3 chromosomes, except one in unclassified contig (ucontig283). Among the 18 duplicated BUSCOs in our assembly, 12 BUSCOs were identified within or between the major three chromosomes.

We compared BUSCOs of our *An. stephensi* assembly with well studied *An. gambiae* reference assembly (fig. S5). A total of 3178 single-copy BUSCOs were common in *An. stephensi* and *An. gambiae*. There were 68 single-copy BUSCOs identified in *An. stephensi* were duplicated in *An. gambiae*, while 18 single-copy BUSCOs in *An. gambiae* were duplicated in *An. stephensi*. Totally, 12 (13499at7147, 19449at7147, 20465at7147, 22971at7147, 31040at7147, 40537at7147, 49088at7147, 57159at7147, 59859at7147, 60890at7147, 62774at7147, 70323at7147) and 5 (24608at7147, 45582at7147, 70842at7147, 74791at7147, 78504at7147) single-copy

BUSCOs were unique to *An. stephensi* and *An. gamabie*, respectively. One duplicated BUSCO (33706at7147) was unique to *An. gambiae*.

#### Immune-related genes/proteins

Using curated sets of immune-related proteins [12], a total of 361 immune genes (31, chrX; 159, chr2; 142, chr3; 11, alternate haplotypes; 18, unclassified contigs) transcribing 394 putative immunity transcripts/proteins that were belonging to 27 gene families were identified in adult *An. stephensi* mosquitoes (table S8). The list of these genes are also available on GitHub (see Data and materials availability). None of them were identified from putative Y-linked contigs. Expansion of many protein families (AMP, APHAG, CLIP, CTL, ML, SCR, SRPN, SRRP, IMDPATH and TOLLPATH) relative to *An. gambiae* account for the large *An. stephensi* immunity-related gene repertoire (fig. S8; table S8). Among 394 proteins, 221 (orthologous to Agam, 205; Aaeg, 8; Cpip, 7; Dmel, 1) were identified as single-copy orthogroup proteins (fig. S8). Out of the remaining 173 proteins, 51 proteins are in a one-to-many relationship, 97 proteins are in many-to-one relationships, and 25 proteins are in a many-to-many relationship (possibly due to gene duplication events) with the known immune proteins. Interestingly, a total of 16 proteins in CLIP, ML, PRDX, SRPN and SRRP families have also been identified to share orthologous proteins with distantly related two mosquitoes Aaeg and Cpip, and Dmel. Our findings showed that the majority of the (i.e. 93% of single-gene) immune proteins were found to share orthologous proteins with *An. gambaie*. Protein expansion in signal modulation IMDPATH family was possibly due to the presence of gram-negative bacteria including symbiont *S. marcescens* [13] (table S8). It also indicated that the rate of gene duplication was higher in *An. stephensi* than in *An. gambiae*. Among 394 immune-related transcripts the top three most abundant transcripts represent signal modulation CLIPs and SRRPs, and recognition CTLs (72, 46 and 30 respectively). Gene losses were observed in families FREPs, GALEs and PPOs. Further studies required on examining the functional importance of specific immune gene family expansions or gene losses in *An. stephensi.* It can facilitate determining particular aspects of immune reactions and evolution to accommodate and reject the pathogens and for its biology including vector competence.

#### Supplementary tables

table S1. Comparison of assembly statistics for *An. stephensi* older and new assemblies.

| Features | Old assembly *(9)* | This assembly |
|---|---|---|
| Total length (bp) | 221,309,404 | 250,632,892 |
| Contig number | 31,761 | 566 |
| Contig N50 | 36,511 | 38,117,870 |
| Scaffold number | 23,371 | 560^ |
| Scaffold N50 | 1,591,355 | 88,747,609* |
| L50 | 40 | 2 |
| GC content (%) | 44.8 | 44.91 |

^Except three major chromosomes, we kept others as contigs; *Scaffold N50 is the length of chr3

table S2. Coordinates of TE sequences that were not found in the existing draft assembly of *An. stephensi* [10]. (separate file)

table S3. Coordinates of exonic TE sequences that were not found in the existing draft assembly of *An. stephensi*. (separate file)

table S4. Coordinates of polymorphic TE sequences that are present in the scaffolds assigned to chromosomes but absent in the alternate haplotype sequences. (separate file)

table S5. Coordinates of putative Y-linked genes supported by multiple uniquely mapping Iso-Seq reads. (separate file) table S6. Genes that are in the top 1% (>64 fold) category of the up- or down-regulated genes after blood feeding. (separate file)

table S7. PBM up or down regulated genes that are either fragmented and missing repetitive sequences like TEs and tandem copies. (separate file)

table S8. Classification of putative immune-related proteins of *An. stephensi*. Comparison in the number of immune-related proteins among *An. stephensi, Aedes aegypti* (Aae), *Anopheles gambiae* (Agam), *Culex quinquefasciatus* (Cpip) and *Drosophila melanogaster* (Flyb/Dmel). (separate file) table S9. A list of representative 23 mosquito genomes and one fly genome from VectorBase/NCBI are included to create a custom database using Kraken2 for classification of *An. stephensi* whole-genome contigs.

| Species Name | Strain | # Scaffolds/Chromosome |
|---|---|---|
| Aedes aegypti | LVP_AGWG | 3 |
| Aedes albopictus | Foshan | 154782 |
| Anopheles albimanus | ALBI9 | 5 |
| Anopheles arabiensis | DONG5 | 1214 |
| Anopheles atroparvus | EBRO | 582 |
| Anopheles christyi | ACHKN1017 | 30369 |
| Anopheles coluzzii | M | 10521 |
| Anopheles culicifacies | A-37 | 5230 |
| Anopheles darlingi | COARI | 2220 |
| Anopheles dirus | WRAIR2 | 1266 |
| Anopheles epiroticus | epiroticus2 | 2673 |
| Anopheles farauti | FAR1 | 116 |
| Anopheles gambiae | PEST | 5 |
| Anopheles maculatus | maculatus3 | 5556 |
| Anopheles melas | CM1001059_A | 5723 |
| Anopheles merus | MAF | 1027 |
| Anopheles minimus | MINIMUS1 | 678 |
| Anopheles quadriannulatus | QUAD4 | 2823 |
| Anopheles sinensis | China | 8007 |
| Anopheles sinensis | SINENSIS | 3101 |
| Anopheles stephensi | Indian_wild_type | 526 |
| Anopheles stephensi | SDA-500 | 1110 |
| Culex pipiens quinquefasciatus | Johannesburg | 3171 |

| | | |
|---|---|---|
| Drosophila melanogaster | A4 | 7 |

table S10. Details of physical map data used in this study. (separate file)

table S11. SRA accession of the publicly available RNAseq data used in this study.

| Sample type | SRA ID |
|---|---|
| adult female | SRR1851030, SRR1851028, SRR1851027, SRR515307 |
| adult male | SRR1851026, SRR1851024, SRR1851022, SRR515308 |
| female larvae | SRR8156253, SRR8156254, SRR8156255, SRR8156256 |
| 0-1h embryo | SRR7061580, SRR7061576 |
| 2-4h embryo | SRR7061579, SRR7061575 |
| 4-8h embryo | SRR7061578, SRR7061574 |
| 8-12h embryo | SRR7061577, SRR7061573 |

table S12. List of 103 unclassified contigs identified as alternate haplotigs using BUSCO Diptera data set and the software Purge_dups.

#### Supplementary Figures

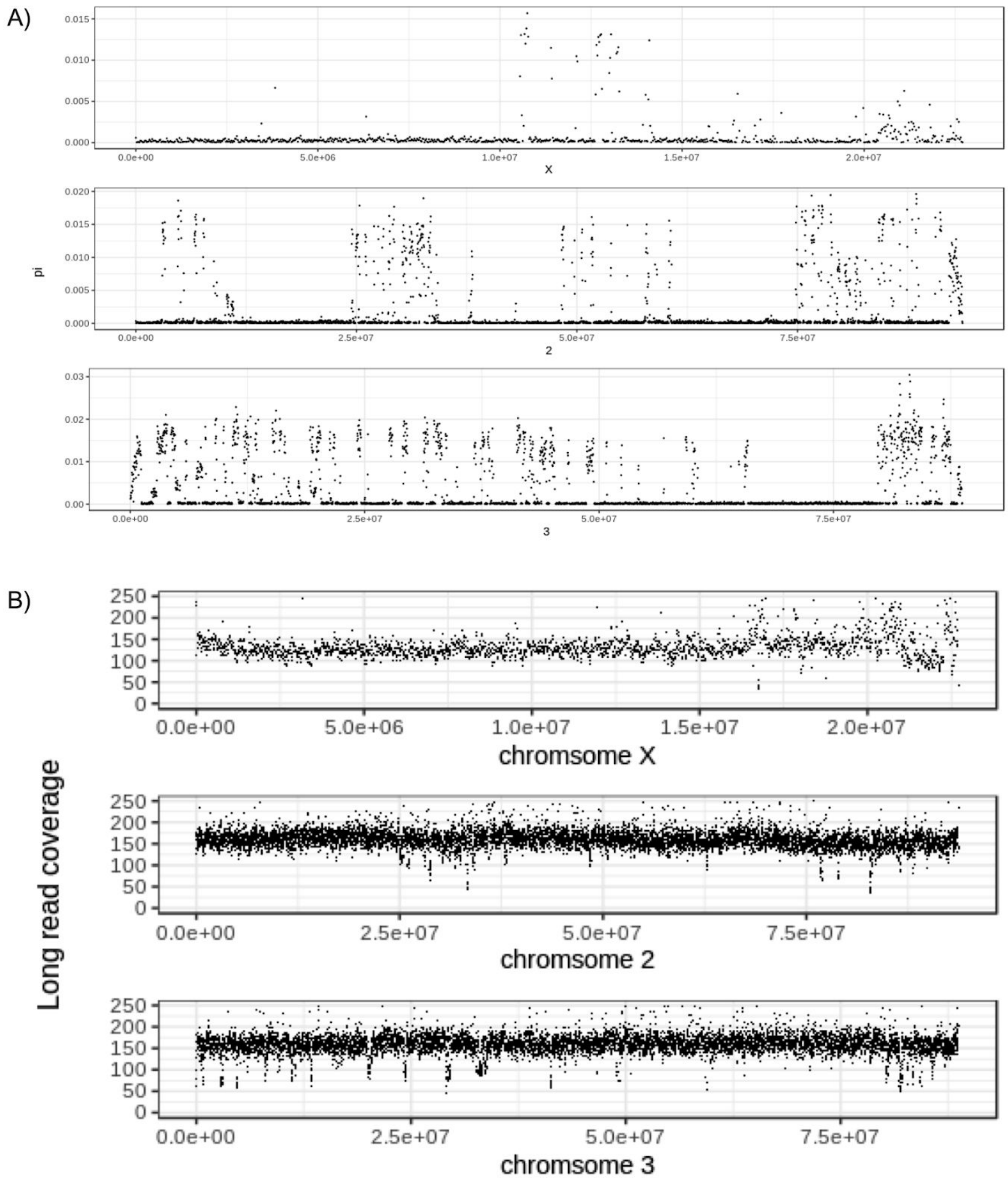


**fig. S1.** (**A**) Heterozygosity across the major chromosome arms (X,2,3) of the inbred sequenced Indian strain of *An. stephensi*. As evidenced here, chromosome 3 has more residual heterozygosity than the other chromosomes. (**B**) Long read coverage in 100bp windows across the scaffolded major chromosome arm sequences. Intermittent coverage drops to non-zero values indicate presence of >1 haplotype in that region. Consistent with the chromosome 3 harboring the highest amount of heterozygosity (refer panel A), such coverage drops are most common in the 3rd chromosome.

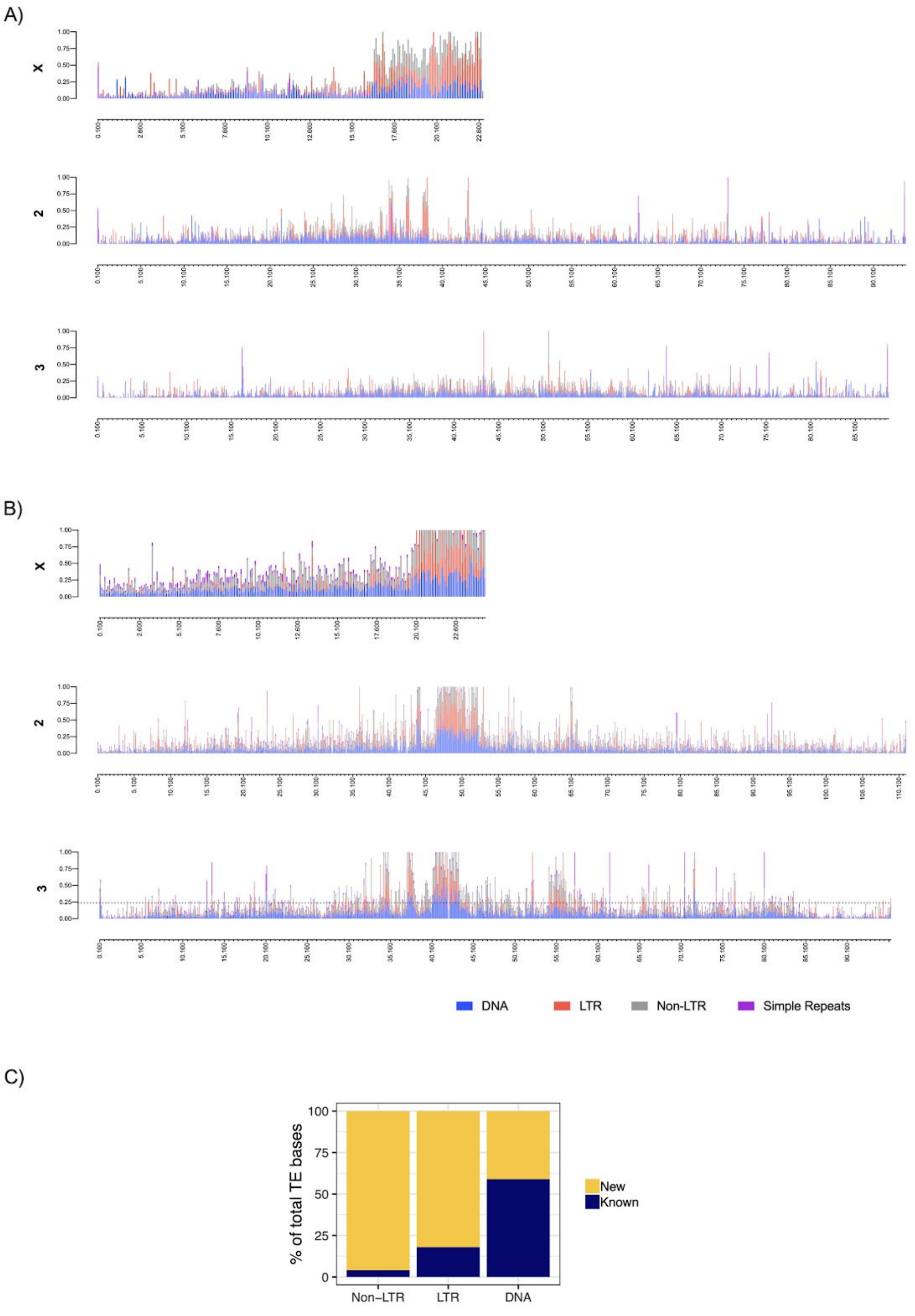


**fig. S2.** The repeat content across the three chromosomes in (**A**) *An. stephensi* and (**B**) *An. gambiae*. The repeat content in the genome was estimated using RepeatMasker and Tandem Repeat Finder. Each bar represents the proportion of different repeat types in 100 Kb non-overlapping windows indicate that the density of repeats on the sex chromosome X is more than that of the autosomes. (**C**) Proportion of TEs (counted in bp) that are present in the new *An. stephensi* assembly but fragmented or absent (new) in the draft assembly [10].

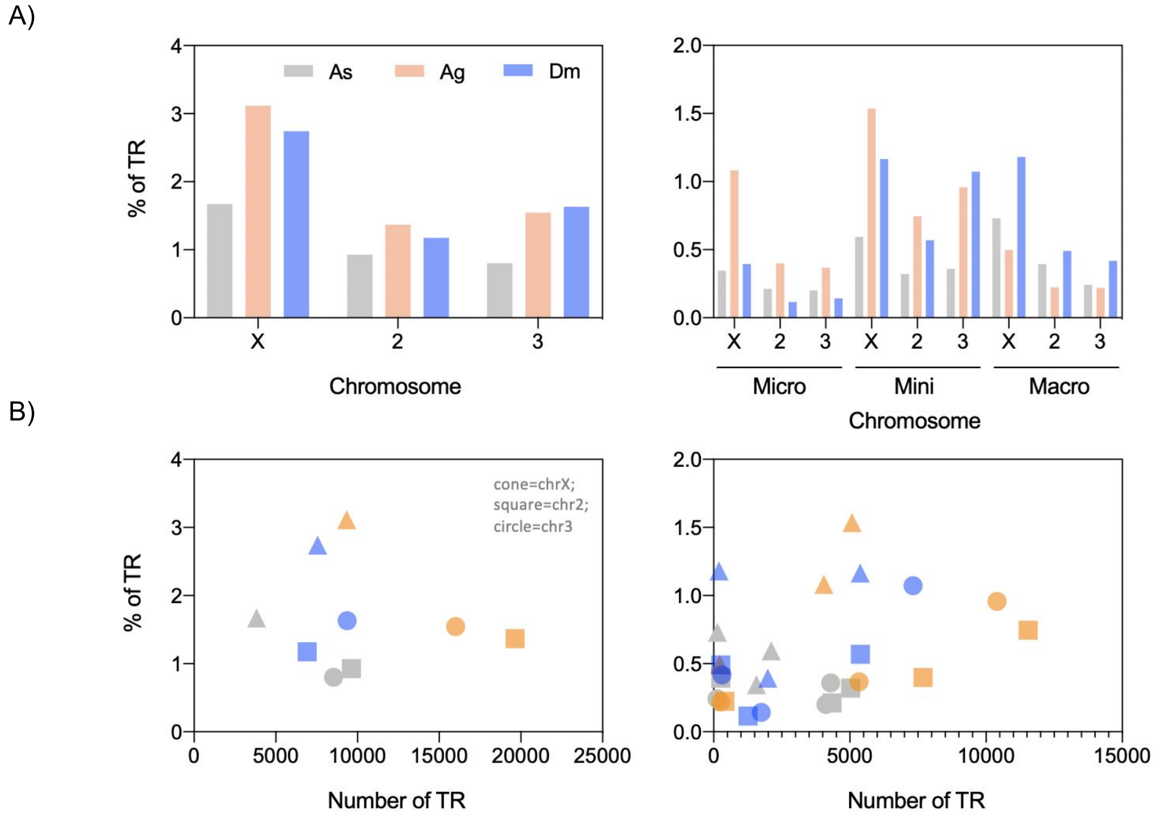


**fig. S3.** (**A**) A comparison of the simple tandem repeats (TR) abundance in three chromosomes of *An. stephensi* (As, grey), *An. gambiae* (Ag, orange) and *D. melanogaster* (Dm, blue) identified using Tandem Repeats Finder. (**B**) The actual number of repeats scattered against the proportion of TR (normalized length of TR). TR consists of a combination of the micro, mini and macrosatellites that were shown in the right panel of the A and B. The proportion of the simple repeats is higher in *An. gambiae* than in *An. stephensi* and *D. melanogaster*.

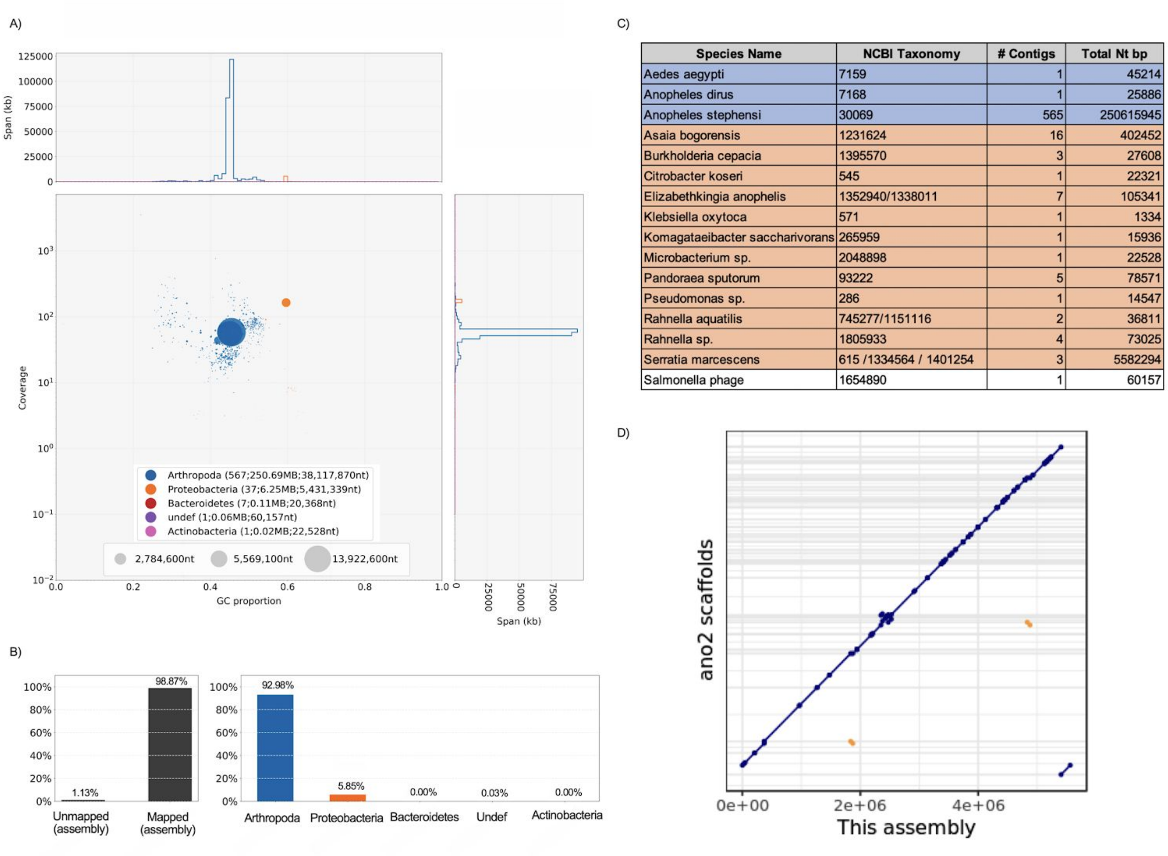


| Species Name | NCBI Taxonomy | # Contigs | Total Nt bp |
|---|---|---|---|
| Aedes aegypti | 7159 | 1 | 45214 |
| Anopheles dirus | 7168 | 1 | 25886 |
| Anopheles stephensi | 30069 | 565 | 250615945 |
| Asaia bogorensis | 1231624 | 16 | 402452 |
| Burkholderia cepacia | 1395570 | 3 | 27608 |
| Citrobacter koseri | 545 | 1 | 22321 |
| Elizabethkingia anophelis | 1352940/1338011 | 7 | 105341 |
| Klebsiella oxytoca | 571 | 1 | 1334 |
| Komagataeibacter saccharivorans | 265959 | 1 | 15936 |
| Microbacterium sp. | 2048898 | 1 | 22528 |
| Pandoraea sputorum | 93222 | 5 | 78571 |
| Pseudomonas sp. | 286 | 1 | 14547 |
| Rahnella aquatilis | 745277/1151116 | 2 | 36811 |
| Rahnella sp. | 1805933 | 4 | 73025 |
| Serratia marcescens | 615 /1334564 / 1401254 | 3 | 5582294 |
| Salmonella phage | 1654890 | 1 | 60157 |

**fig. S4.** Taxonomy classification of contigs from whole-genome assembly of *An. stephensi* using BlobTools. (**A**) Blobplot shows base coverage in a read set of whole-genome sequencing against GC content for contigs. Contigs are colored by phylum with Arthropoda (blue), Proteobacteria (orange), Bacteroides (red) and Actinobacteria (pink). A single contig classified as a DNA virus (purple). Histograms show the distribution of contig length sum along each axis. (**B**) Proportion of classified contigs. (**C**) A total of 92.98% of 613 contigs are classified as Arthropoda, while the remaining 46 are microbial contigs (6.4 Mb) that belong to 12 bacteria and one DNA phage virus. (**D**) Dot plot between *Serratia marcescens* assembly from this study (X axis) and the most contiguous strain of *An. stephensi S. marcescens* (ano2) from NCBI. As evident from the plot, the new assembly has the entire *S. marcescens* genome in a single contig, whereas ano2 has 77 scaffolds. Notably, several structural differences exist between ano2 and new reference strain.

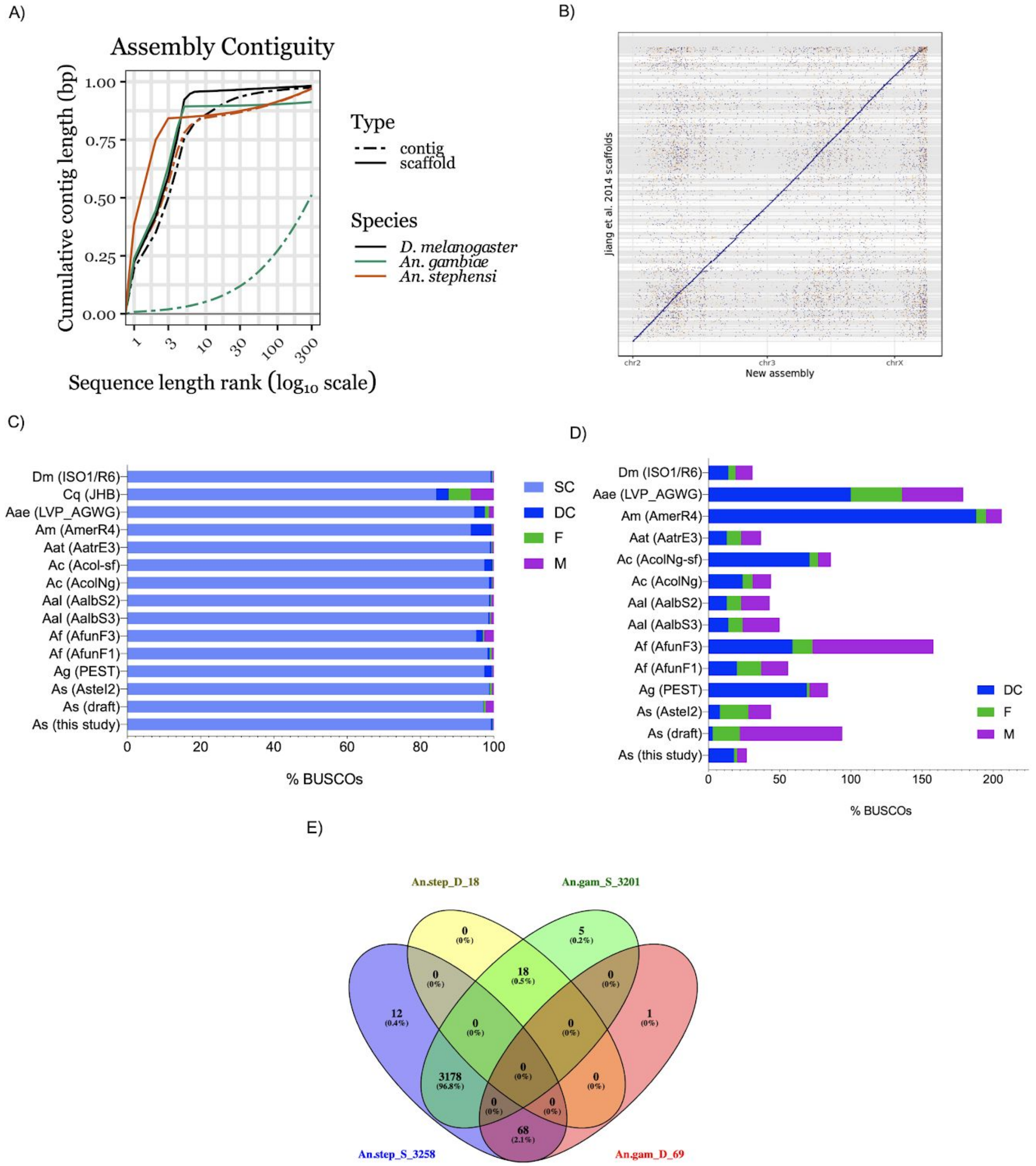

**fig. S5.** (**A**) Comparison of assembly contiguities between *An. stephensi*, *An. gambiae*, and *D. melanogaster* reference assemblies. (**B**) Dot plot between the new reference assembly of *An. stephensi* and the older draft quality assembly [10], Each horizontal line denotes scaffold boundary from the older assembly and each vertical line denotes scaffold boundaries of the assembly from this study. The diagonal demonstrates overall concordance between the two assemblies. The presence of densely positioned numerous horizontal lines (appearing as grey shaded rectangles) demonstrates fragmentation of the older assembly. (**C**) Diptera lineage Benchmarking Universal Single-Copy Orthologs (BUSCO; n=3285) assessment was used to quantify completeness for *An. stephensi* (As; new, draft and Astel2 assemblies), along with the latest version of published chromosome-level assemblies of *An. gambiae* (Ag), *An. funestus* (Af), *An. albimanus* (Aal)*, An. coluzzii* (Ac), *An. atroparvus* (Aat), *An. merus* (Am), *Aedes aegypti* (Aae), *Culex quinquefasciatus* (Cq), and *Drosophila melanogaster* (Dm) genomes [14–17]. It showed that *An. stephensi* new assembly is a best characterized genome among the sequenced malaria vectors. Bar charts show proportions classified as complete C - complete (SC, Single-copy complete; DC, Duplicated complete), F - fragmented and M - missing. (**D**) Comparison of the number of duplicated (D), fragmented (F) and missing (M) BUSCOs among the species (except Cq) shown in A. (**E**) Comparison of BUSCOs of the new *An. stephensi* (An.step) assembly and *An. gambiae* (An.gam). Singleton (An.step_S; An.gam_S) and Duplicated (An.step_D; An.gam_D) BUSCOs were compared to identify common and unique BUSCOs. The number of BUSCOs identified in both species under two categories was also labelled in the venn diagram.

A)

B)

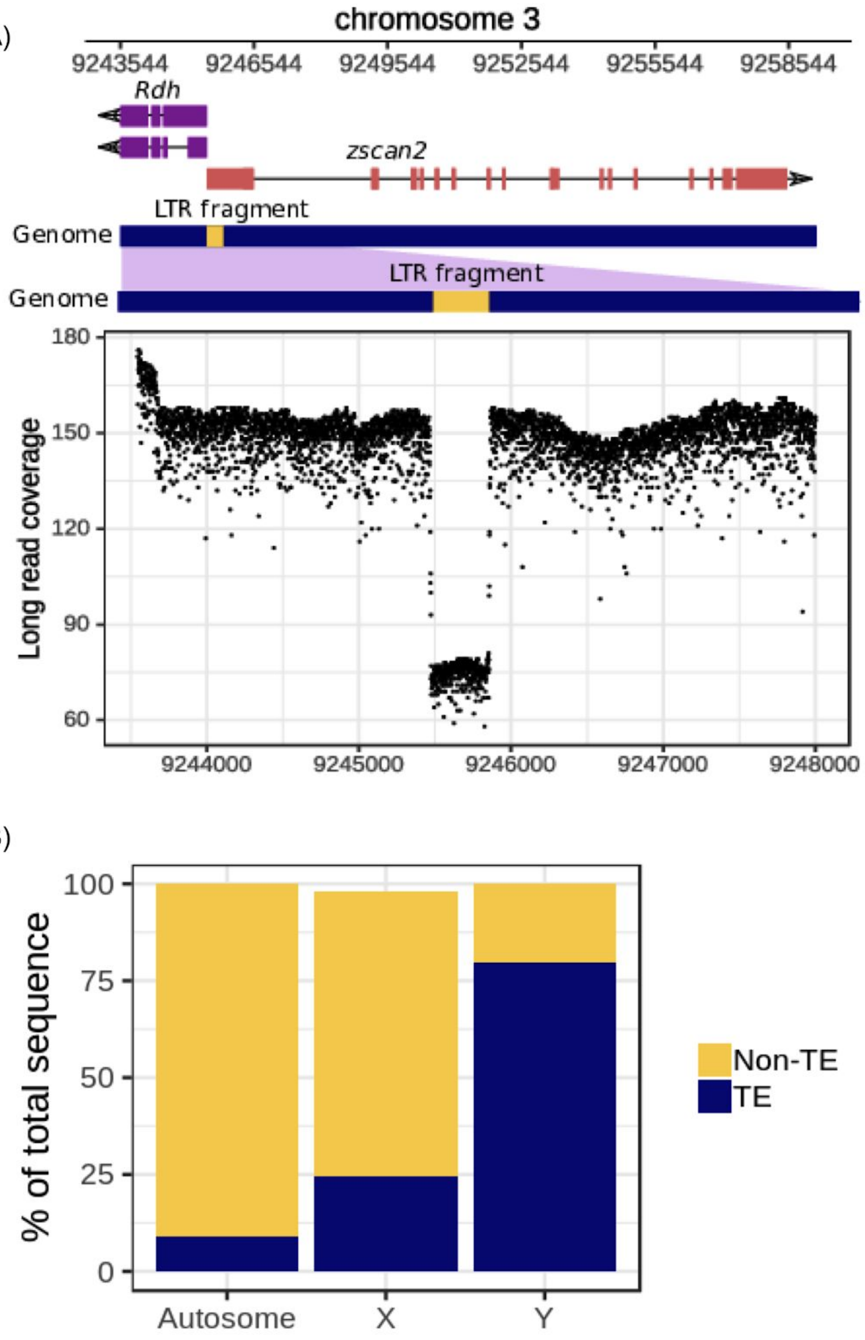


**fig. S6.** (**A**) Polymorphic insertion of a LTR TE fragment within the first exon of *zscan2* and immediately upstream of retinaldehyde dehydrogenase (*Rdh*). The coverage drop to nearly 50% (coverage ~75) for the LTR fragment suggests presence of the insertion in only half of the 3rd chromosomes segregating in the strain. Given that promoters and cis-regulatory sites are often located immediately upstream of a gene, this polymorphic

TE insertion could influence transcription of these two genes. (**B**) Proportion of TE bases in assembled sequences of autosomes (2nd and 3rd), X, and Y. X has more TEs than autosomes but Y has the greatest proportion, with 78% of the Y sequences being TEs.

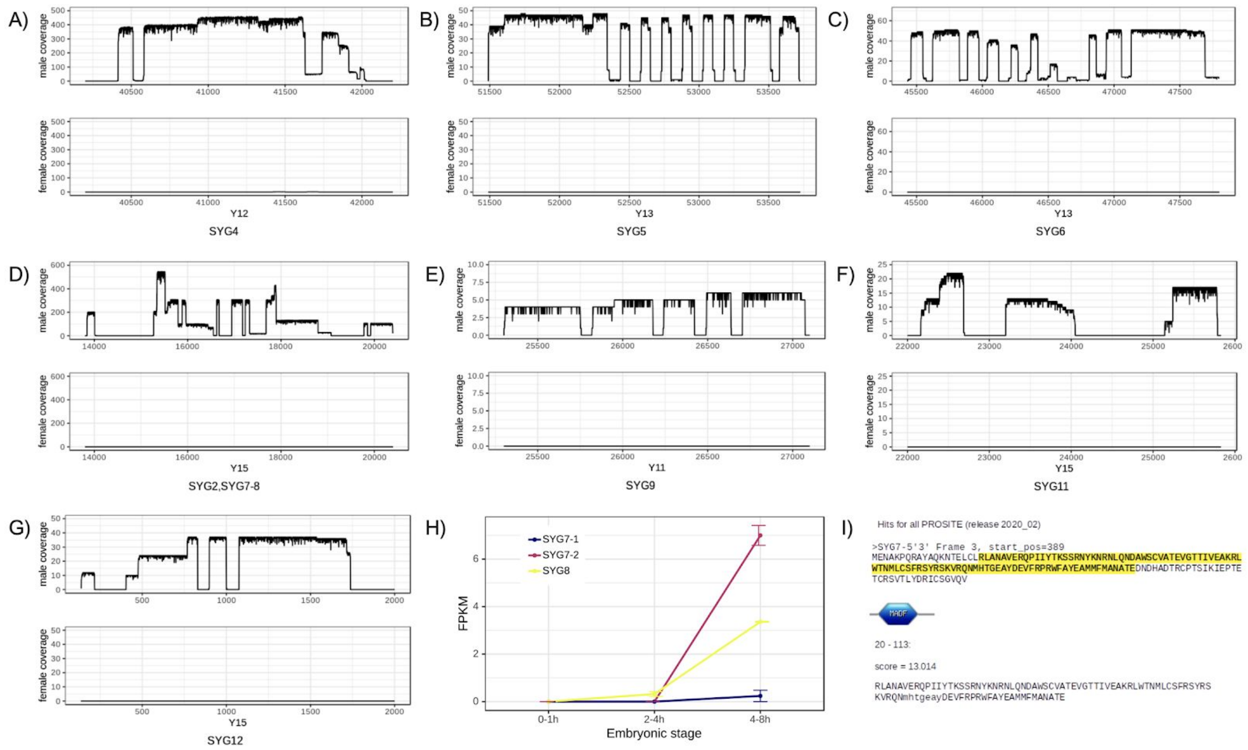


**fig. S7.** Supporting evidence for Y-linked gene (**A**) SYG4, (**B**) SYG5, (**C**) SYG6, (**D**) SYG2, SYG7, SYG8 (detailed gene models of SYG2, SYG7, and SYG8 are depicted in Fig. 1F), (**E**) SYG9, (**F**) SYG11 (the two levels of coverages in the first exon of the gene are due to two transcript isoforms; see table S5), and (**G**) SYG12 (due to the partial fragmentation of the full length mRNA, the 5' end of the transcript (left side of the plot) has lower coverage than the 3' end) from Isoseq read coverage. Exons have more or less uniform coverage from Isoseq reads collected from adult male mRNA, whereas introns are represented by large coverage drops. Consistent with the Y-linkage of this gene, no Isoseq read from adult females map to it. (**H**) Expression of *SYG7* and *SYG8* in early embryos, where both begin to be expressed after 4 hours. (**I**) Presence of MADF domain in the translated protein sequence from SYG7 transcript. The transcript sequence predicted by Iso-Seq reads were translated with an expasy protein translation tool and then scanned with PROSITE.

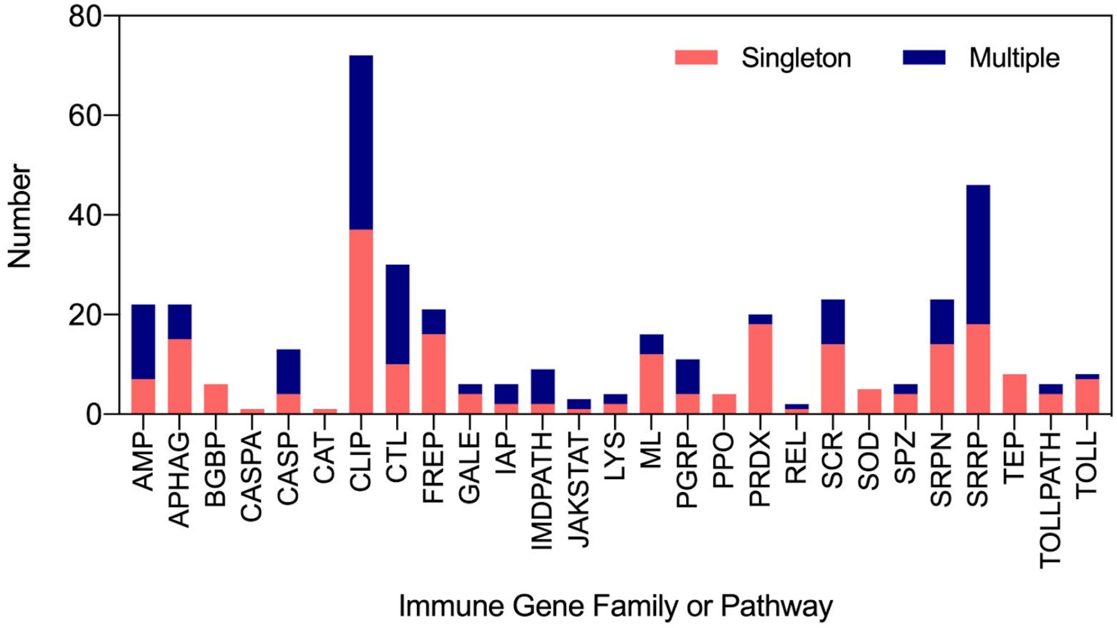


**fig. S8.** The repertoire of putative immune proteins of *An. stephensi* that belong to 27 gene families. Among 394 proteins, 221 are identified as single-copy (orange) while remaining 173 proteins (blue) are identified to have one-to-many, many-to-one and many-to-many (blue) relationships with the curated proteins from the immune database (see table S8).

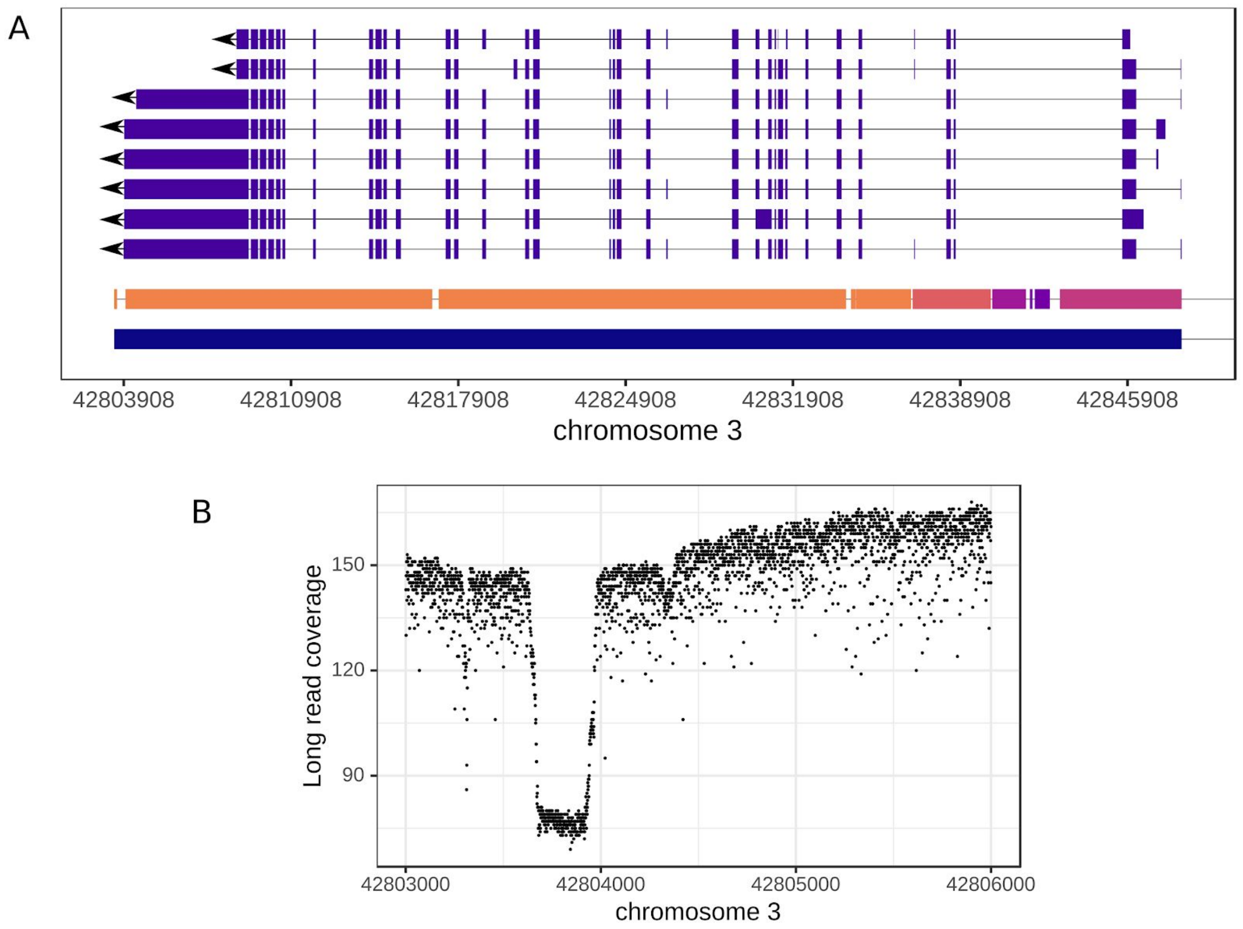


**fig. S9.** kdr gene of *An. stephensi* and presence of SV near the gene; ace2 sequence in the older and new assembly. (**A**) Multiple transcript isoforms of *kdr* easily detected using RNAseq reads mapped to the new *An. stephensi* reference assembly (solid blue bar). However, the older draft assembly has the *kdr* gene split over 6 contigs (different colors of bars above the blue solid bar represents contigs in the older assembly). (**B**) A polymorphic indel immediately downstream of the *kdr* gene, providing evidence that SVs are segregating in this candidate insecticide resistance gene in this strain. Evidence of the indel can be seen as the leftmost gap in the older assembly in A.

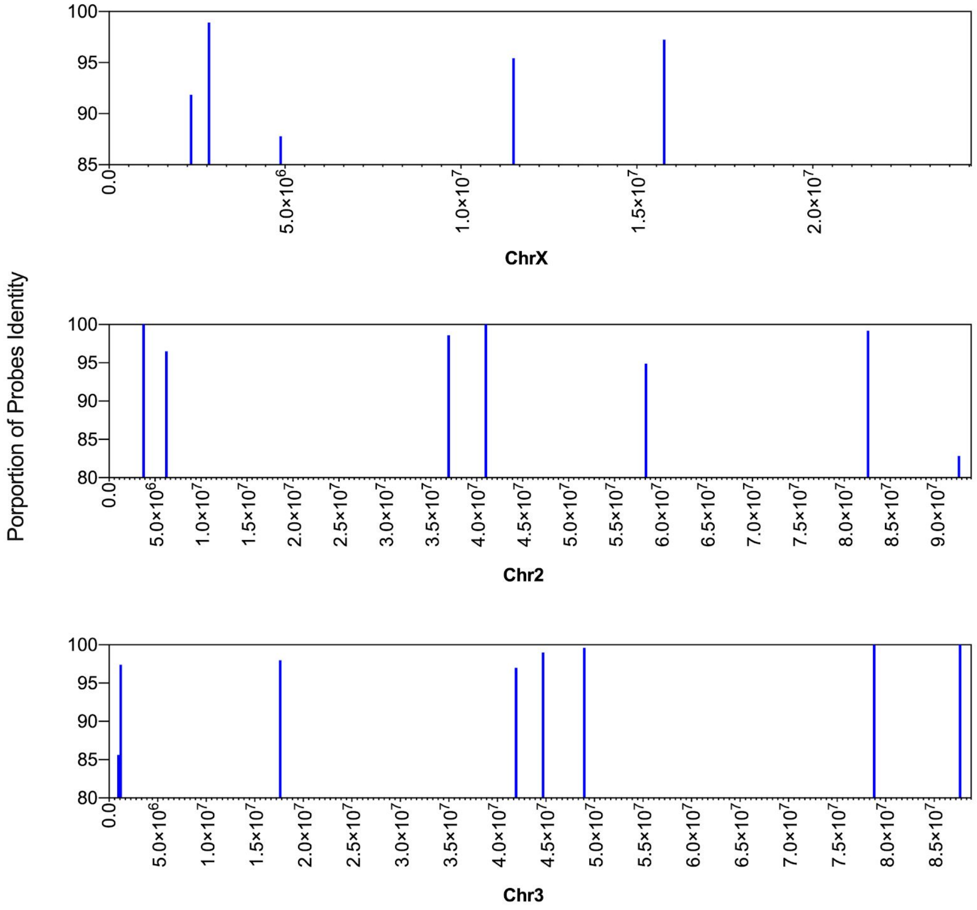


**fig. S10.** Position of 20 physical map probes data against their sequence identity to the new *An. stephensi* genome assembly. The order and orientation of the three chromosomes are examined by MUMmer alignment of 20 gene/probe physical map data (chrX, 5 probes; chr2, 7; chr3, 8) generated from FISH on polytene chromosomes (table S10) [10] against Hi-C chromosome assemblies.

A)

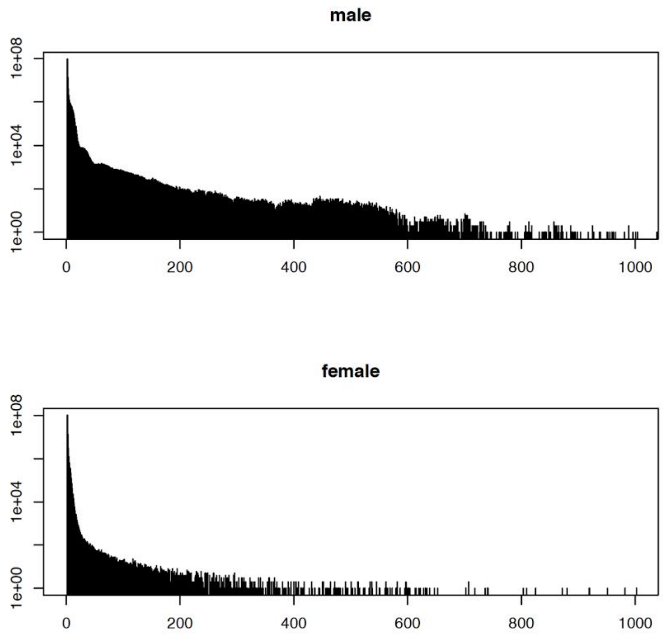


B)

| | | | | | | | | | |
|---|---|---|---|---|---|---|---|---|---|
| chrY15 | maker | mRNA | 15336 | 19330 | + | ANSTEP-UCI_TRAN_00014237-RB | Parent=ANSTEP-UCI_GENE_00014237 | Ma5_7d | Protein of unknown function |
| chrY22 | maker | mRNA | 149789 | 150373 | + | ANSTEP-UCI_TRAN_00013933-RA | Parent=ANSTEP-UCI_GENE_00013933 | | Protein of unknown function |
| chrY28 | maker | mRNA | 6145 | 19836 | - | ANSTEP-UCI_TRAN_00014965-RA | Parent=ANSTEP-UCI_GENE_00014965 | | Similar to Bbx: HMG box transcription factor BBX (Mus musculus OX%3D10090) |

C) D)

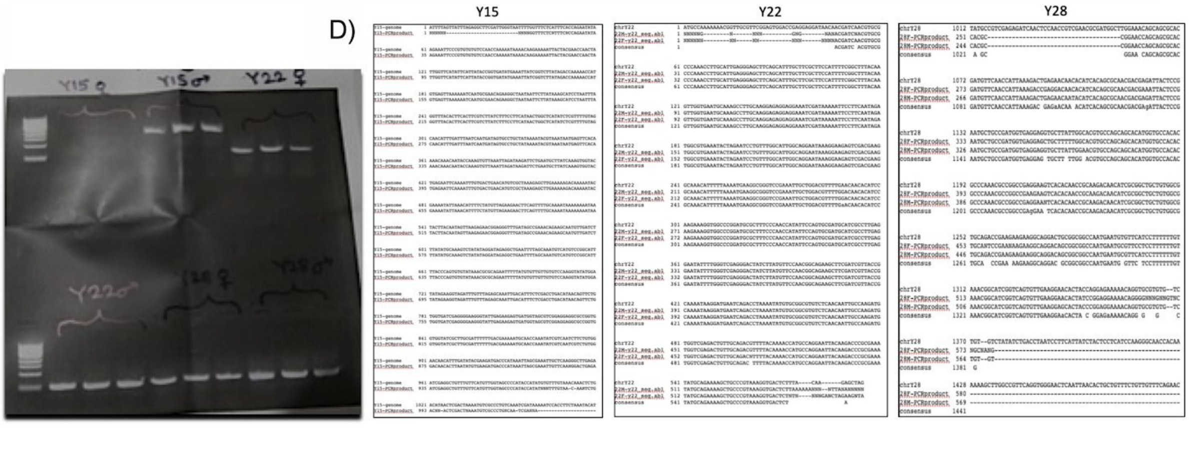


**fig. S11.** (**A**) Male and female specific k-mers identified from the paired-end Illumina reads that were generated from the male (top) and female (bottom) genomic DNA, respectively. (**B**) Experimental validation of putative Y chromosome contigs. Three genomic regions of putative Y contigs that were examined with PCR. These regions also correspond to three predicted genes in MAKER2 (see the full annotation on the

GitHub page or in the genome browser link provided under Data and materials availability). (**C**) Unique sequences were amplified and their PCR products were visualized in Agarose gel. PCR products were purified from the gel and were Sanger sequenced. (**D**) Alignment of Sanger sequenced amplified products and their sequences in the new assembly (and gel picture) confirm male specificity of these contigs.

### Reference


1. Kumar S, Blaxter ML. Simultaneous genome sequencing of symbionts and their hosts. Symbiosis. 2011;55:119–26.

2. Rani A, Sharma A, Rajagopal R, Adak T, Bhatnagar RK. Bacterial diversity analysis of larvae and adult midgut microflora using culture-dependent and culture-independent methods in lab-reared and field-collected Anopheles stephensi-an Asian malarial vector. BMC Microbiol. 2009;9:96.

3. Salter SJ, Cox MJ, Turek EM, Calus ST, Cookson WO, Moffatt MF, et al. Reagent and laboratory contamination can critically impact sequence-based microbiome analyses. BMC Biol. 2014;12:87.

4. Laurence M, Hatzis C, Brash DE. Common contaminants in next-generation sequencing that hinder discovery of low-abundance microbes. PLoS One. 2014;9:e97876.

5. Chen S, Blom J, Walker ED. Genomic, Physiologic, and Symbiotic Characterization of Serratia marcescens Strains Isolated from the Mosquito Anopheles stephensi. Front Microbiol. 2017;8:1483.

6. Cirimotich CM, Dong Y, Clayton AM, Sandiford SL, Souza-Neto JA, Mulenga M, et al. Natural microbe-mediated refractoriness to Plasmodium infection in Anopheles gambiae. Science. 2011;332:855–8.

7. Boissière A, Tchioffo MT, Bachar D, Abate L, Marie A, Nsango SE, et al. Midgut microbiota of the malaria mosquito vector Anopheles gambiae and interactions with Plasmodium falciparum infection. PLoS Pathog. 2012;8:e1002742.

8. Osei-Poku J, Mbogo CM, Palmer WJ, Jiggins FM. Deep sequencing reveals extensive variation in the gut microbiota of wild mosquitoes from Kenya. Mol Ecol. 2012;21:5138–50.

9. Karpe YA, Kanade GD, Pingale KD, Arankalle VA, Banerjee K. Genomic characterization of Salmonella bacteriophages isolated from India. Virus Genes. 2016;52:117–26.

10. Jiang X, Peery A, Hall AB, Sharma A, Chen X-G, Waterhouse RM, et al. Genome analysis of a major urban malaria vector mosquito, Anopheles stephensi. Genome Biol. 2014;15:459.

11. Waterhouse RM, Seppey M, Simão FA, Manni M, Ioannidis P, Klioutchnikov G, et al. BUSCO applications from quality assessments to gene prediction and

phylogenomics. Mol Biol Evol. 2017. doi:10.1093/molbev/msx319.

12. Waterhouse RM, Kriventseva EV, Meister S, Xi Z, Alvarez KS, Bartholomay LC, et al. Evolutionary dynamics of immune-related genes and pathways in disease-vector mosquitoes. Science. 2007;316:1738–43.

13. Nehme NT, Liégeois S, Kele B, Giammarinaro P, Pradel E, Hoffmann JA, et al. A model of bacterial intestinal infections in Drosophila melanogaster. PLoS Pathog. 2007;3:e173.

14. Ghurye J, Koren S, Small ST, Redmond S, Howell P, Phillippy AM, et al. A chromosome-scale assembly of the major African malaria vector Anopheles funestus. GigaScience. 2019;8. doi:10.1093/gigascience/giz063.

15. Kingan SB, Heaton H, Cudini J, Lambert CC, Baybayan P, Galvin BD, et al. A High-Quality De novo Genome Assembly from a Single Mosquito Using PacBio Sequencing. Genes . 2019;10. doi:10.3390/genes10010062.

16. Compton A, Liang J, Chen C, Lukyanchikova V, Qi Y, Potters M, et al. The Beginning of the End: A Chromosomal Assembly of the New World Malaria Mosquito Ends with a Novel Telomere. G3 . 2020;10:3811–9.

17. Lukyanchikova V, Nuriddinov M, Belokopytova P, Liang J, Reijnders MJMF, Ruzzante L, et al. Anopheles mosquitoes revealed new principles of 3D genome organization in insects. Genomics. 2020;:4912.
